# Conservation and divergence of vulnerability and responses to stressors between human and mouse astrocytes

**DOI:** 10.1101/2020.04.17.044222

**Authors:** Jiwen Li, Lin Pan, Marlesa I. Godoy, William G. Pembroke, Jessica E. Rexach, Michael C. Condro, Alvaro G. Alvarado, Mineli Harteni, Yen-Wei Chen, Linsey Stiles, Angela Y. Chen, Ina B. Wanner, Xia Yang, Steven A. Goldman, Daniel H. Geschwind, Harley I. Kornblum, Ye Zhang

## Abstract

Human-mouse differences are a major barrier in translational research. Astrocytes play important roles in neurological disorders such as stroke, injury, and neurodegeneration. However, the similarities and differences between human and mouse astrocytes are largely unknown. Combining analyses of acutely purified astrocytes, experiments using serum-free cultures of primary astrocytes, and xenografted chimeric mice, we found extensive conservation in astrocytic gene expression between human and mouse. However, genes involved in defense response and metabolism showed species differences. Human astrocytes exhibited greater susceptibility to oxidative stress than mouse astrocytes, due to differences in mitochondria physiology and detoxification pathways. Mouse astrocytes, but not human astrocytes, activate a molecular program for neural repair under hypoxia. Human astrocytes, but not mouse astrocytes, activate the antigen presentation pathway under inflammatory conditions. These species-dependent properties of astrocytes may contribute to differences between mouse models and human neurological and psychiatric disorders.

## Introduction

Mice are one of the most widely used experimental animals in biomedical research. Mice and humans share many anatomical, physiological, and genetic features. Because of the ease with which mice can be genetically manipulated and the many paradigms that translate well between species, mouse models have contributed tremendously to our understanding of human biology. Humans and mice had their last common ancestor about 90 million years ago (Springer and Murphy, 2007). Since then, the two species diverged in their body size, life span, ecological niche, behavior, and pathogenic challenges. Therefore, mouse models of human diseases have limitations. For example, many mouse models of neurodegenerative disorders, such as Alzheimer’s disease, amyotrophic lateral sclerosis, and Parkinson’s disease, exhibit milder neuron degeneration phenotypes compared to human patients (Arnold et al., 2013; Bezard et al., 2013; Masliah et al., 2000; Sasaguri et al., 2017). Mouse models of ischemic stroke can often achieve full functional recovery (Ito et al., 2018; Manwani et al., 2011) whereas human patients frequently have irreversible functional deficits. Over 90% of neurological drug candidates with promising animal data fail human clinical trials (Hay et al., 2014; Minnerup et al., 2012), representing a key barrier in translational research. Therefore, identifying cellular and molecular differences between human and mouse brains is urgently needed. The existence of such differences does not invalidate the mouse as an experimental system, but rather underscores the critical need to delineate interspecies difference to optimize their contribution to human disease research. Since access to human cells and tissue for research is limited and many techniques routinely used in studies of mouse models cannot be applied to human patients or samples, mouse models will continue to be an irreplaceable tool of choice for research. The identification of human-mouse differences will accelerate translational research from mouse models to human patients.

Astrocytes constitute at least a third of all cells in the human brain and are critical for the development and function of the central nervous system (CNS) (Anderson et al., 2016; Chung et al., 2013, 2015; Molofsky et al., 2012; Pfrieger and Barres, 1997; Stogsdill et al., 2017; Ullian et al., 2001; Yu et al., 2018), including promotion of neuron survival and growth (Molofsky et al., 2014), stimulation of synapse formation and function (Allen et al., 2012; Blanco-Suarez et al., 2018; Eroglu et al., 2009; Farhy-Tselnicker et al., 2017; Huang et al., 2004; Ma et al., 2016; Parpura et al., 1994; Pascual et al., 2005; Risher et al., 2018; Singh et al., 2016; Stogsdill et al., 2017; Ullian et al., 2001), synapse engulfment (Chung et al., 2013; Tasdemir-Yilmaz and Freeman, 2014; Vainchtein et al., 2018), modulating neuronal activities (Nedergaard, 1994), neural transmitter uptake, ion homeostasis (Kelley et al., 2018a), regulation of vascular function, and metabolite clearance (Xie et al., 2013). Reactive astrogliosis is a spectrum of molecular, cellular, and functional changes in astrocytes that occur in response to a wide range of CNS injuries and diseases, including traumatic injury, stroke, inflammation, epilepsy, brain tumors, Alzheimer’s disease, Parkinson’s disease, and Huntington’s disease (Adams and Gallo, 2018; Bordey et al., 2001; Chaboub et al., 2016; Chen et al., 2008; Gandal et al., 2018; Laug et al., 2019; Liddelow and Barres, 2017; Robel et al., 2015; Sofroniew, 2014). Reactive astrocytes exhibit hypertrophy and upregulation of intermediate filament proteins, change their metabolism (Arneson et al., 2018), secrete pro- and anti-inflammatory cytokines and chemokines that regulate immune cells, produce growth factors that promote neuronal survival and axonal growth, and in some cases form scars that physically limit the spread of injury. Blocking the induction of reactive astrogliosis in mice leads to larger injured regions, increased neuronal death, and worse functional recovery in many scenarios (Anderson et al., 2016; Bush et al., 1999; Herrmann et al., 2008). However, pro-inflammatory mediators can drive astrocytes towards cytotoxic phenotypes (Liddelow et al., 2017). Therefore, reactive astrocytes have both beneficial and harmful roles in the pathogenesis of injuries and diseases.

Reactive astrocytes have been detected in human CNS in numerous conditions such as Alzheimer’s disease, Huntington’s disease, traumatic brain injury, and stroke. Further, astrocyte genes are up-regulated in multiple neuropsychiatric conditions, including bipolar disorder, autism spectrum disorder and schizophrenia, and appear to have some characteristics of reactive astrocytes (Gandal et al., 2018), but have not been extensively characterized. Despite its clear relevance to disease, the biology of reactive astrocytes in humans remains unclear as most of our knowledge is based on studies using murine astrocytes (Krencik et al., 2015, 2017a, 2017b; Majo et al., 2020). Aside from human astrocytes being larger and morphologically more complex than mouse astrocytes (Kelley et al., 2018b; Oberheim et al., 2006, 2009; Oberheim Bush and Nedergaard, 2017), little is known about the similarities and differences between human and mouse astrocytes, especially their responses to disease-relevant perturbations. This knowledge gap has made it challenging to harness knowledge gained from mouse astrocytes to understand the biology of human astrocytes and their roles in neurological disorders.

The lack of understanding of human astrocytes is partly due to technical difficulties in obtaining these cells. Recently, we developed a method to acutely purify human astrocytes and to culture them in serum-free conditions (Zhang et al., 2016). In this study, we used our new method and systematically examined human astrocytes in three conditions: acutely purified, serum-free cultures, and xenografted into mouse brains. We found extensive conservation between human and mouse astrocyte transcriptomes. However, we also found important differences between mouse and human astrocytes that were maintained across all three conditions, including in astrocyte xenografts into mouse brain; demonstrating clear, reproducible differences in the intrinsic transcriptomic properties of human and mouse astrocytes. As suggested by these species-specific differences in gene expression, we identified multiple phenotypic differences in the cellular and molecular responses of human and mouse astrocytes to oxidative stress, hypoxia, inflammatory cytokine treatment, and simulated viral infections. This includes striking differences between human and mouse astrocytes in cell survival, mitochondria physiology, and molecular responses under these disease-relevant perturbations. These findings provide mechanistic understanding of the differences between human and mouse models of neurodegeneration and stroke and uncover new approaches to improve the models.

## Results

### Immunopanned human astrocytes exhibit transcriptome profiles similar to resting astrocytes

The most widely used methods to purify astrocytes were developed 40 years ago, and use serum to select and expand astrocytes (McCarthy and de Vellis, 1980). *In vivo*, serum components only contact astrocytes after injury or disease with blood brain barrier breakdown and induce reactive astrogliosis (Foo et al., 2011; Zamanian et al., 2012). Therefore, serum-selected astrocytes have reactive characteristics (Foo et al., 2011; Zamanian et al., 2012), complicating studies of disease and injury responses. To overcome these complications, we developed an immunopanning method for the acute purification of human astrocytes and a serum-free chemically defined medium that keeps human astrocytes healthy for at least six weeks *in vitro* (Figure 1A-C) (Zhang et al., 2016). Here, we tested whether immunopanned human astrocytes resemble resting or reactive astrocytes by RNA-seq. To evaluate the reactivity status of immunopanned human astrocytes, we assessed the expression of genes previously found to be induced by stroke and inflammation in mouse astrocytes (Zamanian et al., 2012). We found that the expression of these genes is significantly lower in cultured immunopanned astrocytes than in serum-selected astrocytes (Average fold change (FC) =0.18; FDR=0.032; Figure 1D). Therefore, immunopanned human astrocytes provide an improved platform for investigating astrocytic injury, disease responses and reactive astrogliosis compared to serum-containing astrocyte cultures.

**Figure 1.**
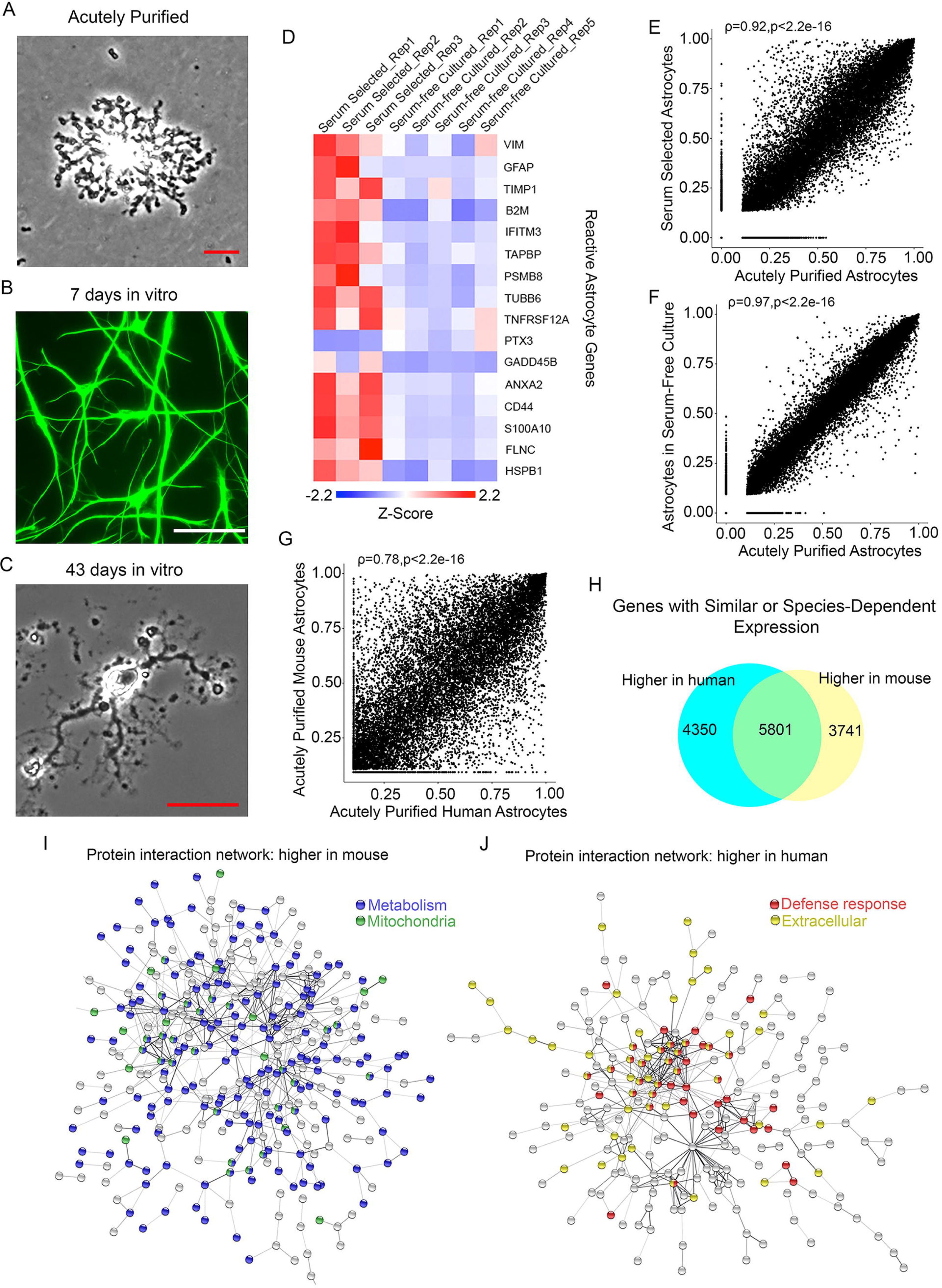
Comparison of astrocyte transcriptomes *in vivo* and *in vitro* and between human and mouse. (A) An astrocyte bound to an anti-HepaCAM antibody-coated petri dish during immunopanning purification. The RNA was extracted immediately after the cells stuck to the dish. These samples were referred to as “acutely purified” thereafter. Scale bar: 10 μm. (B) Astrocytes in serum-free culture stained with anti-GFAP antibodies. Scale bar 50 μm. (C) A bright field image of an astrocyte in serum-free culture. Scale bar: 20 μm. (D) Expression of reactive astrocyte marker genes in serum-selected and serum-free cultures of human astrocytes. Z-score is calculated as (RPKM – average RPKM across all samples) / standard deviation. Genes with FDR<0.1 between serum-selected and serum-free cultures and RPKM>1.5 are shown. (E-F) Scatter plots and Spearman’s correlations of gene expression between cultured and acutely purified human astrocytes using the serum-selected culture method and our serum-free culture method. For each condition, gene expression across 3-5 patient samples was averaged. Only protein coding genes were included. (G) Scatter plot and Spearman’s correlation of gene expression between acutely purified human and mouse astrocytes. Data included children and adult humans as well as developing and adult mice. The age of patients and mice are listed in Supplemental Table 5. (H) Number of genes with similar or species-dependent expression. Genes with percentile ranking in the top two-thirds were included in this analysis to eliminate genes not expressed or expressed at very low levels. Percentile ranking of the expression of each gene were compared across human and mouse astrocyte samples and differences were tested by Welch’s T-test followed by post hoc multiple comparison adjustment using the Benjamini and Hochberg FDR method (Benjamini and Hochberg, 1995). FDR<0.05. (I) Protein interaction networks of genes expressed at higher levels by mouse astrocytes than human astrocytes (Percentile ranking difference >0.4). FDR<0.05. Blue: genes associated with the GO term metabolism. Green: genes associated with the cellular component: mitochondria. (J) Protein interaction networks of genes expressed at higher levels by human astrocytes than mouse astrocytes (Percentile ranking difference >0.4). FDR<0.05. Percentile ranking difference >0.4. Red: genes associated with the GO term defense response. Yellow: genes associated with the cellular component: extracellular.

To examine the extent to which *in vivo* astrocytes could be modeled by immunopanned human astrocytes, we performed immunopanning purification of human astrocytes and harvested RNA (1) immediately after purification to capture the *in vivo* gene signature, as much as possible (referred to as “acutely purified” thereafter) and (2) after 4-6 days of culturing in our serum-free chemically defined medium (referred to as “serum-free cultured” thereafter). We then performed RNA-seq and compared the transcriptomes of acutely purified and cultured human astrocytes. We found that gene expression from serum-free cultures of astrocytes closer resemble acutely purified astrocytes than astrocytes obtained using the traditional serum-selected method (McCarthy and de Vellis, 1980) (Spearman’s correlation = 0.97 vs 0.92, p<0.0001; Figure 1E, F). The expression of the majority of genes (88.1%) are in the same decile in serum free cultures and acutely purified astrocytes (Supplemental Figure 1). Nevertheless, a subset (1.44%) of genes remain differentially expressed between serum-free and “*in vivo*” acutely purified astrocytes (Fold change > 2, false discovery rate (FDR)<0.05, mean Reads Per Kilobase of transcript per Million mapped reads (RPKM) >0.5). For example, acutely purified human astrocytes express higher levels of the glutamate transporter gene Slc1a2 and glutamate receptor gene Grm3 than do serum-free cultured astrocytes (Supplemental Table 1). These genes are likely induced by glutamate released from neurons in the brain environment. Enriched gene ontology (GO) terms in genes differentially expressed between serum-free cultured and acutely purified astrocytes are listed in Supplemental Table 2. Overall, immunopanned human astrocytes recapitulate the expression of the majority of genes expressed by astrocytes *in vivo* and therefore represent an useful platform for studies of human astrocyte biology.

### Human and mouse astrocytes differ in expression of defense response and metabolism genes

Human and mouse diverged about 90 million years ago (Nei et al., 2001; Perlman, 2016). How does adaptation to different ecological niches and body features (different diet, behavior, body size, life span, etc.) influence the physiology of human and mouse brains, and in particular, their astrocytes? To begin to understand the evolutionary differences between human and mouse astrocytes, we compared the acutely purified human astrocytes described above with the corresponding mouse transcriptome data that we previously collected (Zhang et al., 2014, 2016). The overall gene expression profiles showed conservation between human and mouse astrocytes (Spearman’s correlation ρ=0.78; Figure 1G). However, thousands of genes exhibited significant differences in expression between species (8091 genes, FDR<0.05. Figure 1H). To test what genes and pathways exhibit differences between human and mouse astrocytes, we analyzed protein interaction networks and Gene Ontology (GO) using the STRING database web tool for human-mouse differentially expressed genes (Szklarczyk et al., 2019). We found that those genes expressed at higher levels by mouse astrocytes compared to human astrocytes are enriched in multiple GO terms of metabolism (Figure 1I; Supplemental Table 3). In contrast, genes expressed at higher levels by human astrocytes compared to mouse astrocytes are enriched in a single GO term, defense response (Figure 1J; Supplemental Table 3). We analyzed the subcellular localization of proteins encoded by genes differentially expressed between human and mouse astrocytes. Interestingly, mouse astrocytes more highly express genes associated with the compartment “mitochondria” whereas human astrocytes more highly express genes assigned to “extracellular space” (Figure 1I, J), including secreted cytokines.

### The human-specific astrocyte gene signature is intrinsically programmed

The higher expression of defense response genes by human astrocytes could be due to either intrinsic properties or differences in external factors such as other neuronal or glial cell types, systemic or environmental differences between human and mouse samples. To assess whether mouse and human astrocytes are different when exposed to equivalent external environments, we transplanted human astrocytes into a xenograft mouse model and compared them with the neighboring host mouse astrocytes (Benraiss et al., 2016; Han et al., 2013; Osipovitch et al., 2019; Windrem et al., 2008, 2014; Zhang and Barres, 2013). We purified primary human fetal astrocytes, injected them into the brains of neonatal mice (Figure 2A). Here, we used the Rag2 knockout immunodeficient mice that do not have mature lymphocytes to avoid graft rejection. We aged the xenografted chimera mice for about 8 months. We detected widespread distribution of human astrocytes in host mouse brains (Figure 2B-E). We then purified all astrocytes (human and mouse) from the chimeric mice by immunopanning and performed RNA-seq (Methods). We exploited DNA sequence differences between human and mouse genes and were able to separate sequencing reads of human vs. mouse origins at the mapping step (Methods). This approach allowed us to obtain the transcriptome profile of human astrocytes grafted in host mouse brain (Supplemental Table 4).

**Figure 2.**
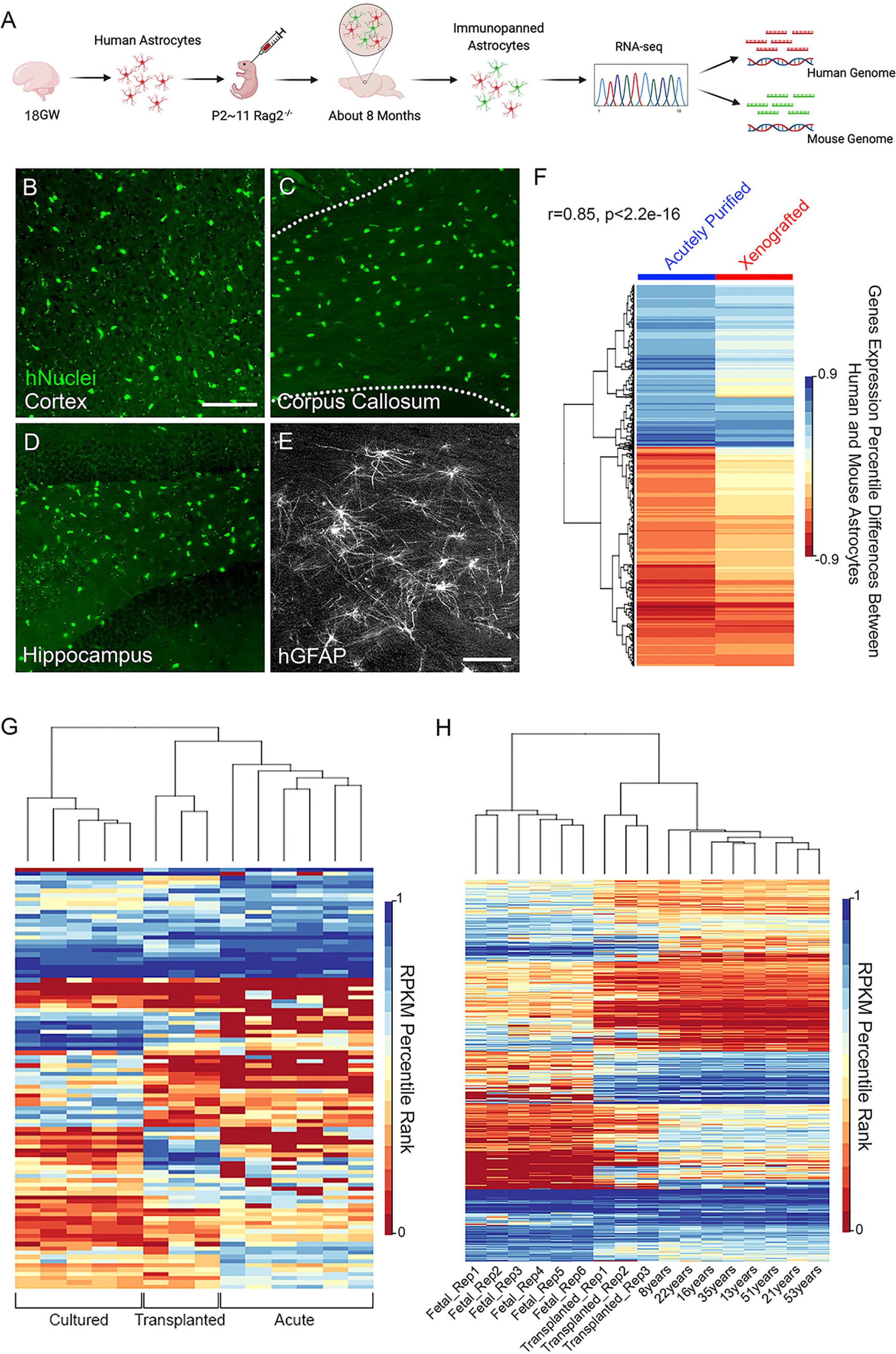
The human-specific astrocyte gene signature is intrinsically programmed. (A) Experimental design. Gestation week 18 primary human astrocytes were purified and injected into the brains of neonatal immunodeficient Rag2 knockout mice. After about 8 months, we purified astrocytes from xenografted mouse brains by immunopanning. These astrocytes from both human grafts and mouse hosts were sequenced together and reads were mapped to human and mouse genomes, respectively. GW, gestational week. P, postnatal day. (B-D) Xenografted human cells in host mouse brains stained with an antibody against human nuclei (green). Scale bar: 100 μm. (E) Xenografted human astrocytes in host mouse brains stained with an anti-GFAP antibody that only reacts with human GFAP. Scale bar: 50 μm. (F) Species differences in gene expression (shown as percentile ranking in human minus percentile ranking in mouse) in xenografted and acutely purified astrocytes highly correlate. Genes with percentile ranking >0.33, species difference FDR <0.05, and species difference in percentile ranking > 0.4 are shown. (G) Non-supervised hierarchical clustering of gene expression of serum-free cultures, acutely purified, and transplanted human astrocytes. Genes with significant differences between cultured and acutely purified astrocytes (FDR<0.05, fold change >4, average RPKM>1) are shown. (H) Non-supervised hierarchical clustering of gene expression of acutely purified astrocytes from patients of different ages and transplanted human astrocytes. Genes with significant differences between age groups (fetal, child, adult. FDR<0.05, fold change >2, maximum RPKM>1) are shown.

To test whether the human-specific astrocyte gene signature is intrinsically programmed or induced by other cell types in the human brain environment, we compared gene expression differences between human and mouse astrocytes using both the acutely purified dataset and xenograft/host dataset. We reasoned that human-mouse astrocyte differences would be attenuated in the xenograft model if the astrocyte differences were driven by human-specific environmental factors. We calculated gene expression differences between human and mouse astrocytes based on the acutely purified dataset (hmDiff_acute) and the chimera dataset (hmDiff_chimera) (Figure 2F). We found a positive correlation between hmDiff_acute and hmDiff_chimera (Pearson’s correlation=0.60). The correlation was even stronger when we only included genes with large species differences (Pearson’s correlation=0.85 for genes with percentile difference >0.4. Figure 2F). We also calculated gene expression correlations between transplanted human astrocytes and acutely purified human/mouse astrocytes and found that transplanted human astrocytes resemble human astrocytes more than mouse astrocytes (Correlation coefficient of transplanted human vs acutely purified human is significantly higher than that of transplanted human vs acutely purified mouse: p<0.0001. Supplemental Figure 2A, B). These analyses suggest that the human-specific astrocyte gene signature is largely intrinsically programmed and environmental influence from neurons and other cell types in the human brain environment only plays minor roles.

We also found that certain astrocytic genes that are expressed in acutely purified astrocytes *in vivo* and are lost in culture, are regained in xenografted astrocytes (Figure 2G). One challenge that has limited human astrocyte research is the difficulty in obtaining mature cells for experimental manipulations. Availability of fresh healthy adult human brain tissue is limited. Stem cell derived human astrocytes mostly resemble developing stages (Krencik et al., 2011; Sloan et al., 2017; Tchieu et al., 2019). We determined that xenografted human astrocytes can reach mature stages that are difficult to access in *in vitro* models (Figure 2H; Supplementary Figure 2C-E); allowing us to further observe persistent of mouse and human transcriptomic differences across a broad developmental range. Therefore, across acutely purified, cultured, and xenografted conditions, we were able to identify consistent species differences in astrocyte transcriptomic profiles. Furthermore, the xenograft model provides a much needed platform for studies of mature human astrocytes *in vivo*.

### Human astrocytes are more susceptible to oxidative stress than mouse astrocytes

The differences in defense response and metabolism between human and mouse astrocytes may either be neutral drifts in evolution that does not affect fitness or have functional significance. It is unclear whether baseline differences may predispose them to respond differently to stressors encountered in disease and injury. Identifying such differential responses may improve interpretation of results from mouse models and guide translational research. To test responses of human and mouse astrocytes, we treated them with several disease-relevant stimuli including oxidative stress, hypoxia, simulated viral infection, and an inflammatory cytokine.

Oxidative stress is produced by reactive oxygen species (ROS) such as peroxides, superoxide, hydroxyl radical, singlet oxygen, and alpha-oxygen. ROS are byproducts of normal metabolism in most cell types in the body (Turrens, 2003). Importantly, in pathogen invasion, tissue damage, and inflammation, immune cells such as neutrophils and macrophages produce high levels of ROS that help fend off pathogen infections but may also damage healthy cells in infected tissue (Yang et al., 2013). In the brain, oxidative stress is a key pathological process underlying neurodegenerative disorders (such as Alzheimer’s disease (Tönnies and Trushina, 2017), Parkinson’s disease (Puspita et al., 2017), Huntington’s disease (Zheng et al., 2018), amyotrophic lateral sclerosis (Bozzo et al., 2017)), stroke (Rodrigo et al., 2013), and traumatic injury (Rodriguez-Rodriguez et al., 2014).

To examine responses of human and mouse astrocytes to oxidative stress, we purified human and mouse astrocytes from developmentally equivalent stages (gestational week 17-20 human brains and P1-3 mouse brains. See Methods for details on the matching of developmental stsages). Astrocytes have been shown to be regionally heterogeneous (Bayraktar et al., 2020; Chai et al., 2017a; Farmer et al., 2016; Glasgow et al., 2014; Hochstim et al., 2008; John Lin et al., 2017; Miller et al., 2019; Tsai et al., 2012; Zhang and Barres, 2010). Therefore, whenever possible, we used matching anatomical locations in human and mouse brains. We used whole cerebral cortex for astrocyte purification for all mouse samples and a subset of human samples with clearly identifiable cerebral cortex. In cases where identification of cerebral cortex was difficult due to tissue fragmentation, we selected, to the best of our knowledge, fragments most likely to be cortex (large thin sheets).

We performed immunopanning purification to obtain human and mouse astrocytes, plated them at similar densities, and cultured them using identical growth media. To examine responses of human and mouse astrocytes to oxidative stress, we treated cell cultured 3 days *in vitro* (div) with 100 μM H_2_O_2_. We then examined cell survival by staining with the live cell dye calcein-AM and the dead cell dye ethidium homo-dimer 18 hours after onset of treatment (Figure 3A). We found that human astrocytes are much more susceptible than mouse astrocytes to oxidative stress. The survival rates were 0.29±0.02 for human astrocytes and 0.54±0.01 for mouse astrocytes (Figure 3B-D. p<0.001. Data represent average±SEM unless otherwise noted). Since plating density, H_2_O_2_ concentration and treatment duration may affect cell survival, we tested a different combination of these conditions and found, again, human astrocytes are much more susceptible than mouse astrocytes to oxidative stress (Supplemental Figure 3).

**Figure 3.**
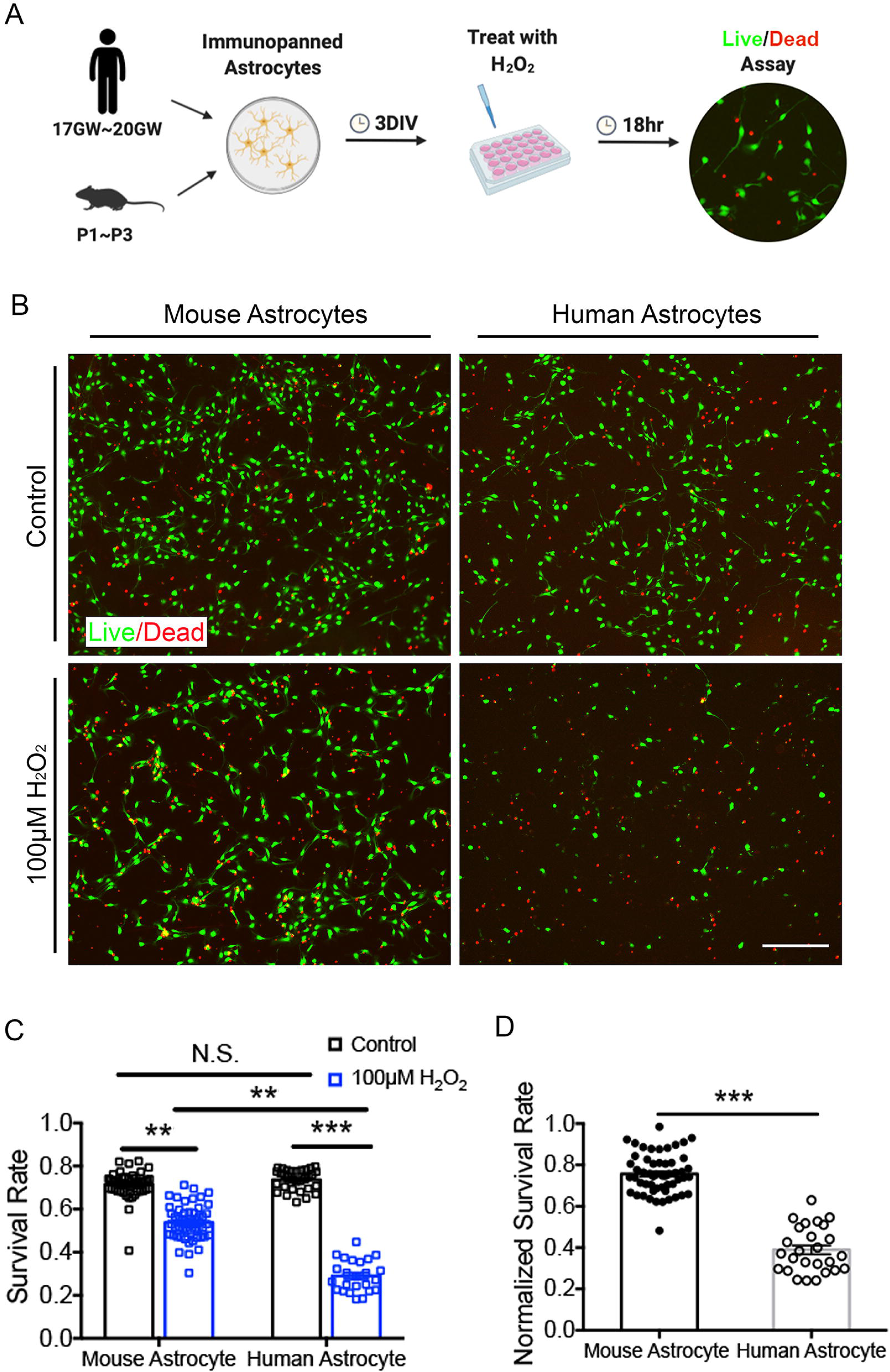
Human astrocytes are more susceptible to oxidative stress than mouse astrocytes. (A) Experimental design. Hr, hour. (B) Human and mouse astrocytes treated with H_2_O_2_ or medium control stained with the live cell dye calcein-AM (green) and the dead cell dye ethidium homodimer (red). Scar bar: 200μm. (C) Survival rate. *, p<0.05. **, p<0.01. ***, p<0.001. N.S., not significant. N=2-5 imaging fields each from 2-4 wells of cultured cells from per patient/litter of mice, generated from 3 patients and 6 litters of mice in C and D. (D) Survival rate of astrocytes treated with H_2_O_2_ normalized to the survival rate of medium control treated cells.

### Mouse astrocytes have faster mitochondria respiration than human astrocytes

Mitochondria are both sources and targets of ROS (Turrens, 2003). The mitochondrial respiration chain produces ROS. On the other hand, ROS (either endogenous or exogenous) can damage mitochondrial function. Furthermore, mitochondria play important roles in cell death (Vakifahmetoglu-Norberg et al., 2017). Therefore, to search for cellular mechanisms underlying the striking difference in susceptibilities of human and mouse astrocytes to oxidative stress, we examined mitochondria metabolism in these cells. We purified human and mouse astrocytes and cultured them under the same conditions. We then used Agilent Seahorse Respirometry to assay mitochondria metabolism.

Although human astrocytes are larger compared to mouse astrocytes *in vivo* (Kelley et al., 2018b; Oberheim et al., 2009) and *in vitro* (Zhang et al., 2016), the basal respiration rate per mouse astrocytes (oxygen consumption rate, OCR) is almost twice as high compared with human astrocytes (mouse: 1.41±0.11 pmol/min per 1000 cells. Human: 0.71±0.09 pmol/min per 1000 cells. P<0.05. Figure 4A). Furthermore, respiration for ATP production is also higher in mouse astrocytes than in human (mouse: 1.02±0.06 pmol/min per 1000 cells. Human 0.58±0.08 pmol/min per 1000 cells. P<0.05. Figure 4B).

**Figure 4.**
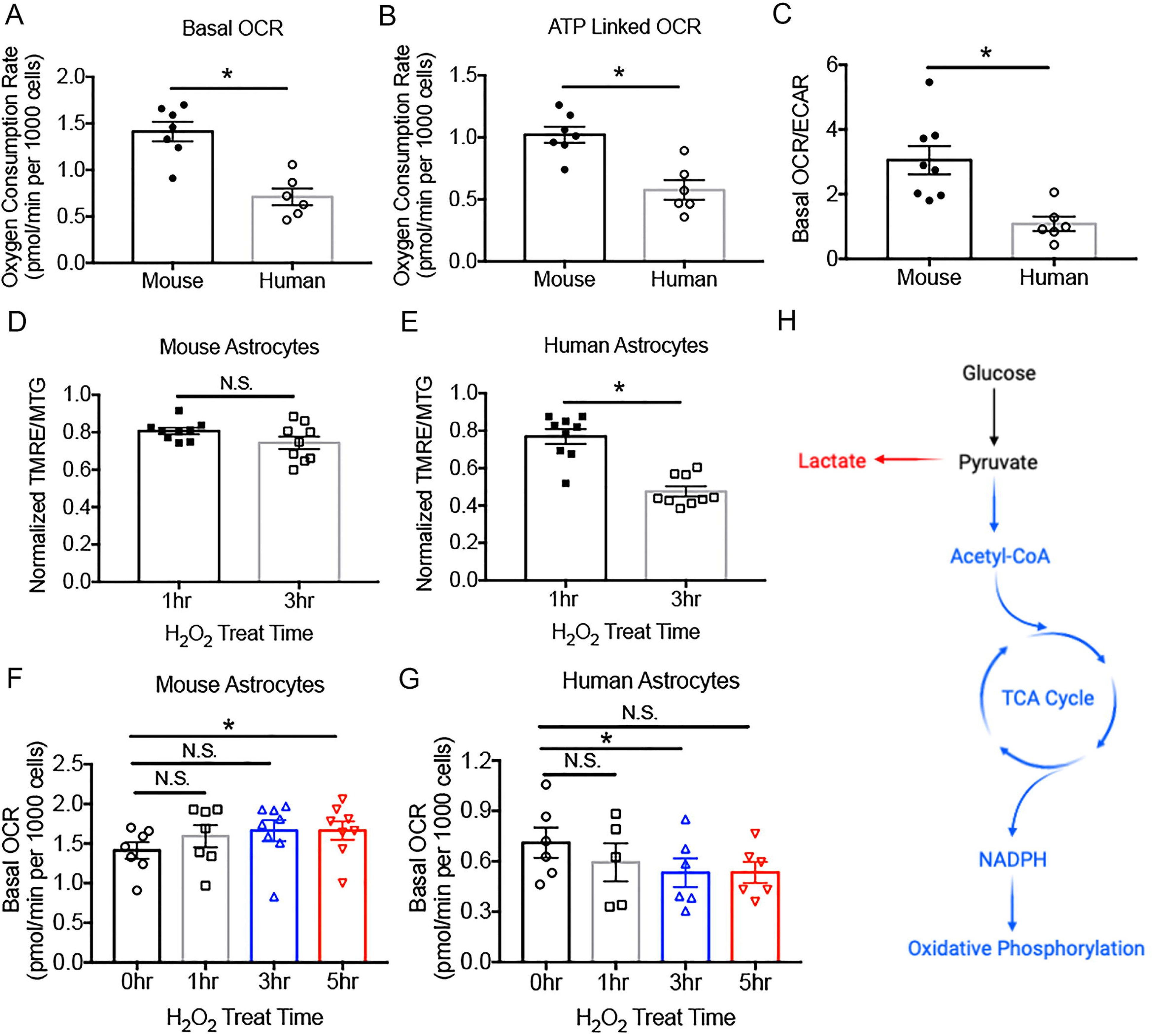
Mitochondria metabolism differences between human and mouse astrocytes. (A) Basal oxygen consumption rate (OCR) of mouse and human astrocytes. Each data point (circle or square) represents 1-3 wells of astrocyte cultures prepared from one human patient or one litter of 8-10 mice throughout the paper. N=1-3 wells of cultured cells per patient/litter of mice, generated from 4 litters of mice and 3 patients in A-C, F-G. (B) OCR linked to ATP production in the presence of oligomycin. (C) The ratio of OCR over extracellular acidification rate (ECAR). (D-E) TMRE fluorescence (mitochondria membrane potential) normalized by MTG (general mitochondria dye) fluorescence. Data shown are H_2_O_2_ treated conditions normalized to medium control treated conditions. N=3 wells of cultured cells per patient/litter of mice, generated from 3 litters of mice and 3 patients. (F-G) Basal OCR of astrocytes treated with 100 μM H_2_O_2_. (H) Diagram of glucose metabolism.

Differences in mitochondria respiration rates in human and mouse astrocytes raised the question of whether energy substrates were utilized differently in these cells. Glucose is the predominant energy substrate in healthy brains (Mergenthaler et al., 2013). Glucose undergoes glycolysis and becomes pyruvate. Pyruvate may go into one of two alternative metabolic pathways (Figure 4H): (1) pyruvate can be converted to acetyl-CoA, go through the tricarboxylic acid (TCA) cycle, and eventually be converted to substrates of oxidative phosphorylation and produce ATP. This process occurs intracellularly within astrocytes. (2) Pyruvate can be converted to lactate and exported to extracellular space. Neurons can take in lactate and use it as an energy substrate, although the astrocyte-neuron-lactate-shuttle hypothesis remains controversial (Yellen, 2018). To examine the usage of glucose by the two alternative pathways in human and mouse astrocytes, we used the Seahorse Respirometry’s pH sensitive electrodes to measure extracellular acidification rate (ECAR), an approximate measure of lactate production and glycolysis rate, and compared ECAR to OCR, an approximate measure of oxidative phosphorylation rate. We found that the OCR/ECAR ratio is higher in mouse astrocytes than in human astrocytes (Figure 4C). These results showed that mouse astrocytes utilize a larger proportion of glucose for oxidative phosphorylation, which provides energy for astrocytes themselves, whereas human astrocytes utilize a larger proportion of glucose for lactate production, which may serve as an energy substrate for neurons (Yellen, 2018).

Having found metabolic differences between human and mouse astrocytes in unperturbed conditions, we next tested changes in mitochondrial metabolism and physiology under oxidative stress in human and mouse astrocytes. We treated the astrocytes with 100 μM H_2_O_2_ and measured OCR 1, 3, and 5 hours after onset of treatment. We found that mouse astrocytes exhibited a small increase whereas human astrocytes exhibited substantial reduction of OCR under oxidative stress (Figure 4F, G. Mouse: 0 hr 1.41±0.11, 5 hr 1.66±0.12, p<0.05. Human: 0 hr 0.71±0.09, 3 hr 0.53±0.08, p<0.05.). Therefore, mitochondria from mouse astrocytes are highly resilient to oxidative damage. They may work harder as an adaptive response to oxidative damage and as a result produce more ATP that cellular protective pathways can use (see the section on detoxification pathway below). In contrast, mitochondria from human astrocytes are very quickly damaged and cannot keep up with cellular energy demand when exposed to oxidative stress.

To further examine the physiological status of mitochondria under oxidative stress, we performed fluorescence imaging using tetramethylrhodamine ethyl ester (TMRE), a dye sensitive to the membrane potential across the inner membrane of mitochondria (Crowley et al., 2016). We found that mitochondria membrane potential remains largely stable in mouse astrocytes but depolarizes quickly in human astrocytes (Figure 4D, E. Mouse 0.81±0.02 at 1hr, 0.74±0.03 at 3hr, not significant. Human 0.77±0.04 at 1hr, 0.48±0.03 at 3hr, p<0.05). Therefore, mitochondria in human astrocytes are more susceptible to oxidative damage compared to mouse astrocytes.

### Mouse astrocytes express higher detoxification pathway genes than human astrocytes

We showed that mouse astrocytes have an oxidative phosphorylation rate twice as fast as that of human astrocytes. Presumably, more ROS are produced by mouse astrocytes than by human astrocytes. We also showed that mouse astrocytes are more resilient than human astrocytes to oxidative stress. How do we reconcile these observations? One hypothesis is that in mouse astrocytes, adaptive mechanisms, such as more efficient detoxification pathways, have evolved under high ROS conditions and renders protection against oxidative stress. To test this hypothesis, we examined the function of peroxisome, an organelle involved in detoxification of ROS (Reddy and Mannaerts, 1994). We blocked mitochondria oxidation with Antimycin A, which binds and inactivates Complex III (Ma et al., 2011). Remaining non-mitochondria oxygen consumption has a large contribution from peroxisomal oxidation (Reddy and Mannaerts, 1994). We found that non-mitochondria oxygen consumption rate is higher in mouse astrocytes than in human astrocytes (Figure 5C), consistent with the possibility that peroxisome detoxification operates faster in mouse astrocytes than human astrocytes. To evaluate molecular differences in ROS detoxification pathways, we purified human and mouse astrocytes by immunopanning and performed RNA-seq immediately after purification (Zhang et al., 2014, 2016) and after 5-6 days of culture in serum-free conditions (this study). We found that the gene encoding a major peroxisomal ROS detoxification enzyme, catalase (Dringen et al., 2005), is expressed at 3-6 fold higher levels by mouse astrocytes than by human astrocytes (RPKM: acutely purified, 2.04±0.24 for human astrocytes and 5.67±0.38 for mouse astrocytes. *In vitro*, 1.14±0.10 for human astrocytes and 7.60±0.71 for mouse astrocytes. Figure 5A). An additional molecular pathway, the pentose phosphate pathway, produces NADPH that neutralizes ROS (Patra and Hay, 2014). The rate limiting step of the pentose phosphate pathway is catalyzed by glucose-6-phosphate dehydrogenase (G6PD). By RNA-seq, we found that G6PD gene expression is 2-10 fold higher in mouse astrocytes than in human astrocytes (RPKM: acutely purified, 0.17±0.05 for human astrocytes and 2.14±0.48 for mouse astrocytes. *In vitro*, 2.05±0.32 for human astrocytes and 4.39±0.32 for mouse astrocytes. Figure 5B). We also explored other major detoxification pathways and found generally comparable expression by human and mouse astrocytes. Taken together, a higher rate of peroxisome oxidation and higher amounts of catalase and G6PD may protect mouse astrocytes against oxidative stress, whereas human astrocytes are more susceptible (Figure 5 D, E).

**Figure 5.**
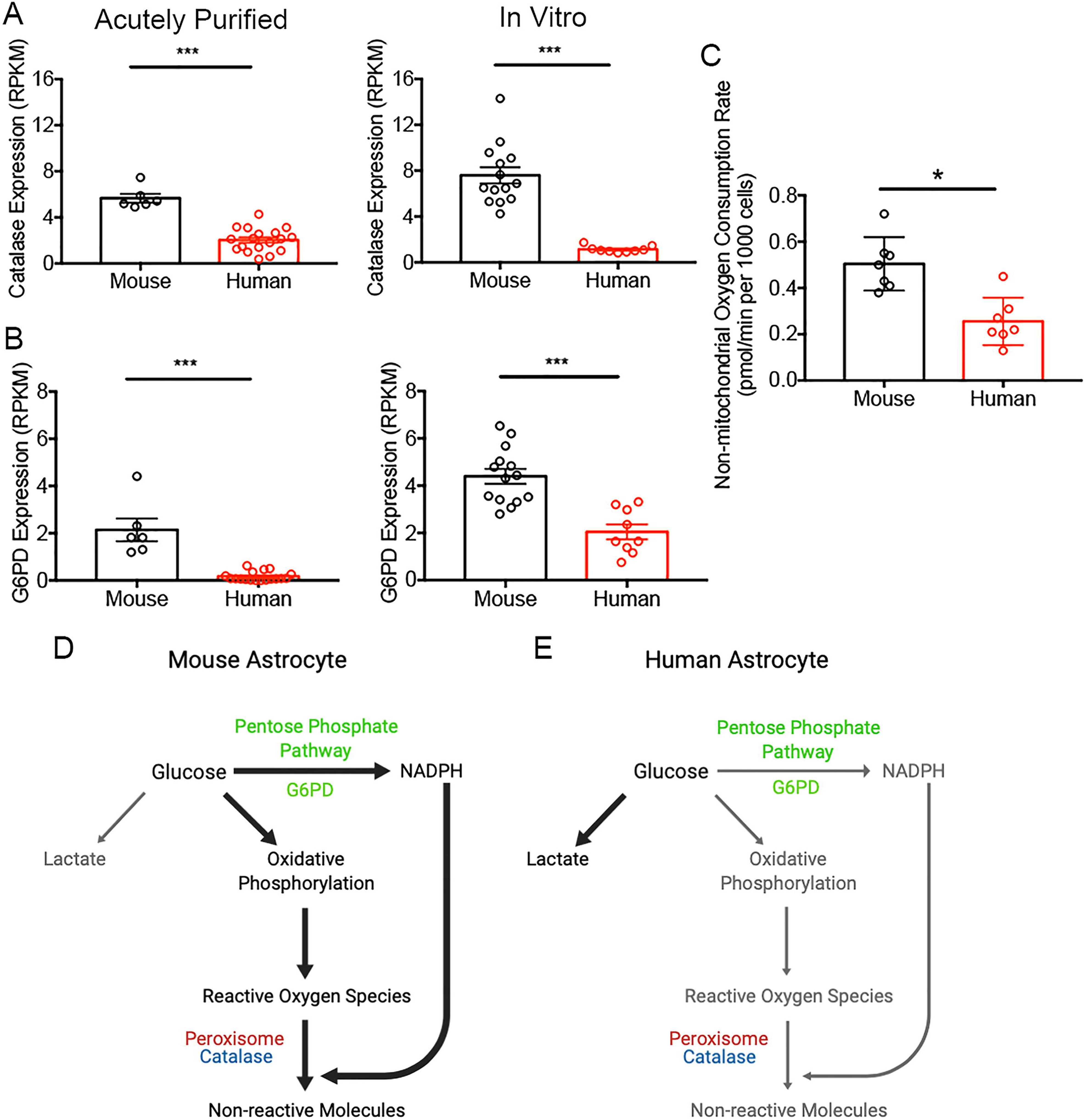
Detoxification pathway differences between human and mouse astrocytes. (A-B) Expression of reactive oxygen species detoxification pathway genes Catalase (A) and G6pd (human, ortholog gene name: G6pdx in mouse) (B) by acutely purified astrocytes and serum-free cultures of astrocytes determined by RNA-seq. n=6 litters of mice and 18 human patients for acutely purified samples. n=14 litters of mice and 9 human patients in vitro. Samples included children and adult patients as well as developing and adult mice (The age of patients and mice is listed in Supplemental Table 5) (C) Non-mitochondrial oxygen consumption rate measured in the presence of antimycin-A. N=7 wells of cultured cells from each species generated from 4 litters of mice and 3 patients. (D-E) A model of glucose metabolism and detoxification pathways in human and mouse astrocytes. The widths of the arrows represent rate of the metabolic processes. Mouse astrocytes have higher rates of oxidative phosphorylation, which presumably produces more reactive oxygen species than human astrocytes. Higher abundance of detoxification pathways genes and higher peroxisomal activity in mouse compared to human may protect the cells against oxidative damage.

Although our *in vitro* functional experiments were all performed using developing astrocytes, we obtained RNA-seq data from adult astrocytes from humans and mice (Supplemental Table 5). Notably, the species differences persisted throughout development and adulthood (Figure 5 A, B). Oxidative stress is a core pathological process in a range of neurological conditions including neurodegenerative disorders such as Alzheimer’s disease (Tönnies and Trushina, 2017), Parkinson’s disease (Puspita et al., 2017), Huntington’s disease (Zheng et al., 2018), and amyotrophic lateral sclerosis (Bozzo et al., 2017). Mouse models of neurodegenerative disorders often have milder phenotypes compared to human patients (Arnold et al., 2013; Bezard et al., 2013; Masliah et al., 2000; Sasaguri et al., 2017). Our findings suggest that differences in astrocytic responses to oxidative stress may be one of the reasons that make mouse models of neurodegeneration more resilient than human patients.

### Hypoxia induces a molecular program for neural growth in mouse but not human astrocytes

Adult mouse models of ischemic stroke often achieve full functional recovery (Ito et al., 2018; Manwani et al., 2011) whereas adult human stroke patients usually have irreversible functional deficits. Dozens of human clinical trials of neural protective drug candidates that improved recovery in mouse models of stroke have all failed so far (Minnerup et al., 2012). Hypoxia is one of the key physical changes in ischemic stroke. Responses of mouse astrocytes to hypoxia have been closely examined previously (Zamanian et al., 2012) but responses of human astrocytes are largely unknown.

We exposed human and mouse astrocytes to hypoxia (Figure 6A). We found that both human and mouse astrocytes exhibited similarly high cell survival and had normal healthy morphology under hypoxia and control conditions. We then performed RNA-seq of all treated and control human and mouse astrocytes. To assess transcriptional responses, we used a combination of differential expression and weighted gene co-expression network analysis (WGCNA) (Methods; Supplemental Figure 4; Supplemental Table 6). Modules identified by WGCNA as significantly up- or down-regulated under treatment application are enriched for genes up- and down-regulated, respectively, under that same condition.

**Figure 6.**
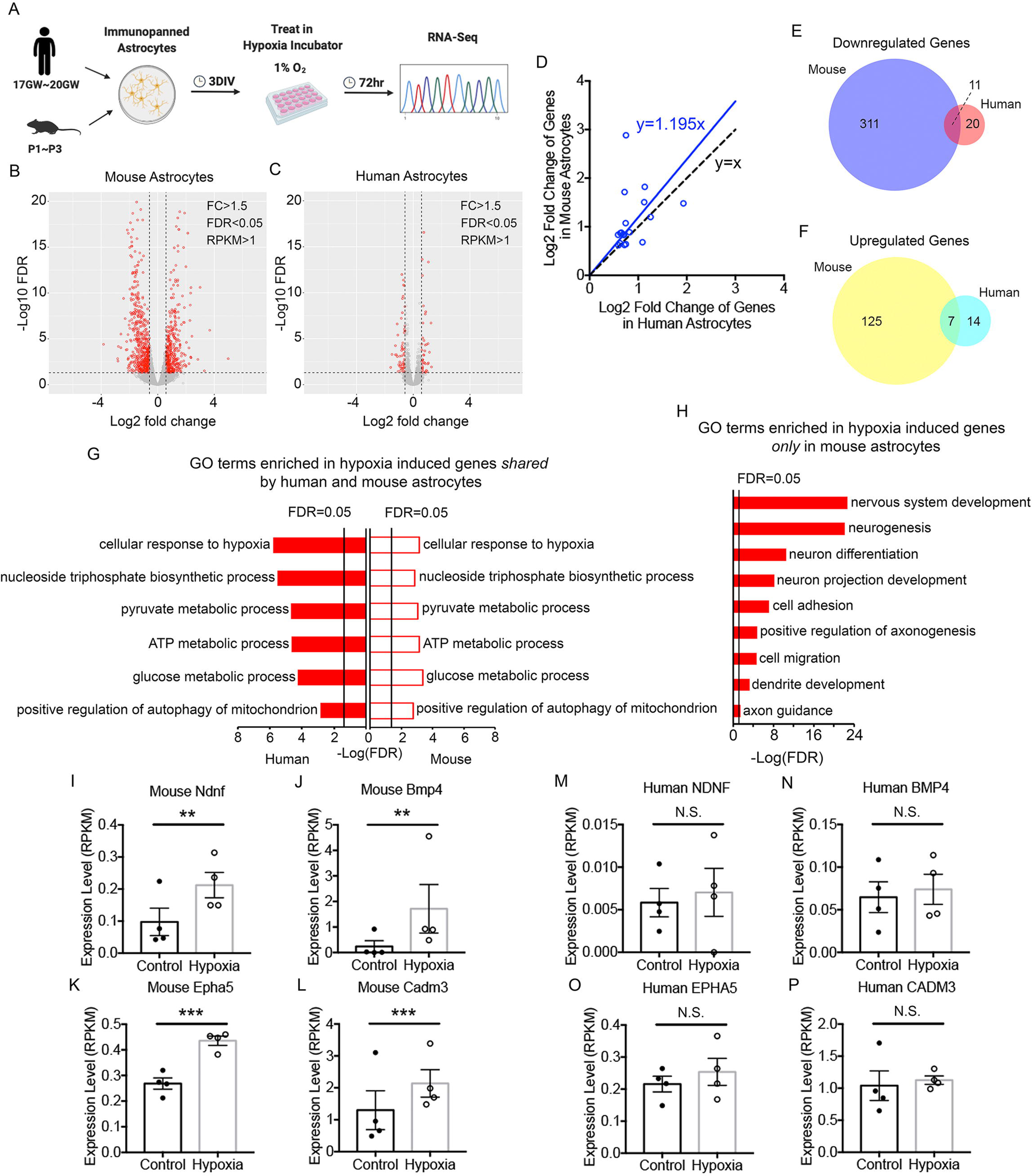
Molecular responses of human and mouse astrocytes to hypoxia. (A) Experimental design. (B-C) Volcano plots of genes significantly different between hypoxia and control conditions. Each red dot represents a significantly different gene. (D) Comparison of fold change of hypoxia-induced genes in human and mouse astrocytes. Genes with FDR<0.05 in both species, FC>1.5, and average RPKM of control/treated >1 are included. Only genes that showed significant change in both species are included. Blue line: linear regression trendline. Black line: predicted trendline of y=x if there were no species differences. (E-F) The number of significantly up- or down-regulated genes in hypoxia treated human and mouse astrocytes. Genes with FDR<0.05, fold change >1.5, and average RPKM of control or treated groups >1 are shown. (G) Top shared gene ontology (GO) terms enriched in hypoxia-induced genes in human and mouse astrocytes ranked by FDR. (H) Development associated GO terms *only* enriched in hypoxia-induced genes in mouse but not in human astrocytes. (I-P) Expression of genes associated with the “nervous system development” GO term in hypoxia and control treated human and mouse astrocytes. n=4 litters of mice and 4 human patients.

When we examined the extent to which hypoxia induced genes are shared between human and mouse astrocytes, we found that 3.4% (11 out of 322) genes downregulated in human astrocytes are also downregulated in mouse astrocytes and 5.3% (7 out of 132) genes upregulated in human astrocytes are also upregulated in mouse astrocytes, demonstrating partial conservation of hypoxia responses between human and mouse (Figure 6E, F. Upregulated overlap: 13.8-fold higher than expected by chance; p=3.65e-07. Downregulated overlap: 6.0-fold higher than expected by chance; p=7.50e-07. See also Supplemental Figure 5). Gene Ontology (GO) terms and KEGG pathway analyses found that, as expected, genes upregulated in both human and mouse astrocytes are enriched in the GO term hypoxia response and the Hypoxia Inducible Factor 1 (HIF1) pathway (Figure 6G and Supplemental Table 3). Interestingly, astrocytes from both species upregulate genes involved in glycolysis and positive regulation of mitochondrial autophagy. Glycolysis provides an alternative pathway to generate energy without oxygen and autophagy of idling mitochondria may conserve resources within cells. Human and mouse astrocytes both downregulate genes involved in amino acid transport (Supplemental Table 3), which is potentially another adaptive change to slow down protein synthesis when energy supply is limited.

Despite the partial conservation of hypoxia responses between human and mouse, hypoxia induced stronger molecular changes in mouse astrocytes relative to human astrocytes in terms of number of differentially expressed genes (454 in mouse vs. 52 in human. >1.5-fold change, FDR <0.05, average RPKM >1. Figure 6B, C, E, F) and effect size (Figure 6D). Genes upregulated by hypoxia in mouse, but not in human astrocytes are enriched in GO terms such as nervous system development, neurogenesis, neuron differentiation, and axon guidance (Figure 6H and Supplemental Table 3) and include the genes which encode Ndnf, a growth factor, Bmp4, a morphogen, Epha5, an axon guidance molecule, and Cadm3, a cell adhesion molecule (Figure 6I-P and Supplemental Table 7, 8). Furthermore, we found a module (grey60) to be upregulated by hypoxia in mouse astrocytes, but not in human astrocytes (Supplemental Table 6 and Supplemental Figure 4). This module is involved in development and cell adhesion, corroborating our finding that hypoxia induces a molecular program that aids neural repair in mouse astrocytes, but not in human astrocytes. These differences may contribute to the differences in functional recovery and responses to drug candidates between human patients and mouse models of stroke.

### Poly I:C and TNF**α** induce antigen presentation pathways in human but not mouse astrocytes

Many viruses, such as human immunodeficiency virus, new world alpha viruses, and some flaviviruses (e.g. Zika virus) are capable of infecting CNS cells, inducing neuroinflammatory responses, and causing acute or long-lasting neurological deficits (Michalicová et al., 2017). Astrocytes, along with microglia, are CNS resident cells that modulate neuroinflammation. However, responses of human astrocytes to viral infections and their consequences to CNS homeostasis and function are poorly understood. In addition to viral infections, neuroinflammation is a core pathological component of a range of neurological conditions such as traumatic injury, stroke, neurodegeneration, and aging. TNFα is one of the major pro-inflammatory cytokines involved in neuroinflammation. TNFα induces reactivity of mouse astrocytes. Although it has been assumed that TNFα would induce similar changes in human astrocytes, the actual effect of this key pro-inflammatory cytokine on human and mouse astrocytes has not been compared.

We exposed human and mouse astrocytes to the viral mimetic double stranded RNA, Poly I:C, or TNFα (Figure 7A). Both human and mouse astrocytes exhibited similarly high cell survival and had normal healthy morphology under treatment and control conditions. We then performed RNA-seq of all treated and control human and mouse astrocytes.

**Figure 7.**
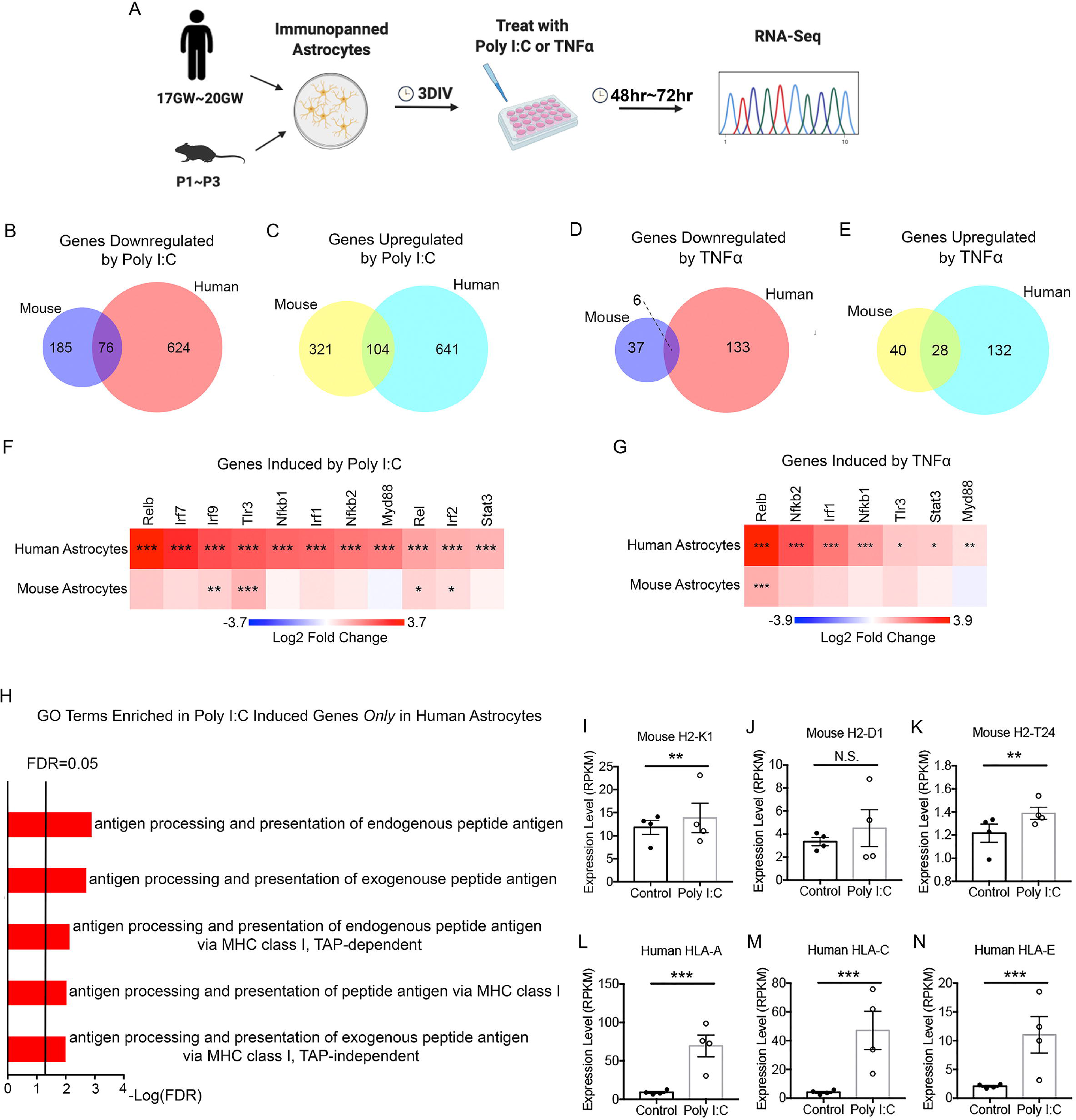
Molecular responses of human and mouse astrocytes to Poly I:C and TNFα. (A) Experimental design. (B-E) The number of significantly up- or down-regulated genes in Poly I:C and TNFα-treated human and mouse astrocytes. Genes with FDR<0.05, fold change >1.5, and average RPKM of control or treated groups >1 are shown. (F, G) Fold change of Tlr3, NFκB, and interferon response pathway genes in Poly I:C and TNFα-treated human and mouse astrocytes. Stars represent significance determined by DESeq2. (H) Selected antigen presentation-related GO terms enriched in Poly I:C-induced genes *only* in human astrocytes. (I-N) Expression of the top 3 highest expressing MHC Class I antigen presentation genes in Poly I:C-treated and control human and mouse astrocytes. n=4 litters of mice and 4 human patients.

In contrast to hypoxia, we found that both Poly I:C and TNFα induced stronger molecular responses in human astrocytes relative to mouse astrocytes in terms of number of differentially expressed genes (Figure 7B-E) and effect size (Supplemental Figure 6). We treated mouse astrocytes with recombinant TNFα proteins from either human or mouse sources and found similar numbers of differentially expressed genes (data not shown). We next examined the extent to which Poly I:C and TNFα-induced gene changes is shared between human and mouse astrocytes. We found a significant proportion (10.9%; 76 out of 700) of downregulated genes in human astrocytes are also downregulated in mouse astrocytes and a significant proportion (14.0%; 104 out of 745) of upregulated genes in human astrocytes are also upregulated in mouse astrocytes under Poly I:C treatment (Figure 7B, C; upregulated: 1.7-fold higher than expected by chance; p=2.79e-07; downregulated: 2.2-fold; p=4.89e-12). A significant proportion (4.3%; 6 out of 139) of genes downregulated by TNFα in human astrocytes are also downregulated in mouse astrocytes and a significant proportion (17.5%; 28 out of 160) genes upregulated by TNFα in human astrocytes are also upregulated in mouse astrocytes, reflecting partial conservation between human and mouse (Figure 7D, E; upregulated: 13.1-fold; p=2.53e-25; downregulated: 5.1-fold; p=9.85e-04). We also observed a significant correlation of fold change in mouse vs fold change in human (Poly I:C, 0.236. TNFα 0.192; Supplemental Figure 5). GO term and KEGG pathway enrichment analysis of the conserved genes (Supplemental Table 3) revealed enrichment for genes involved in response to cytokines and other organisms, consistent with known effect of inflammatory cytokines and viral infections.

Furthermore, we examined genes induced by Poly I:C or TNFα only in human astrocytes. These genes are enriched in GO terms include antigen processing and presentation of peptide antigen via Major Histocompatibility Complex (MHC) class I (Figure 7H). The expression of the three highest expressing MHC Class I genes in human astrocytes (HLA-A, HLA-C, and HLA-E) and mouse astrocytes (H2-K1, H2-D1, and H2-T24) is shown in Figure 7I-N. These genes showed modest or no increase in Poly I:C or TNFα-treated mouse astrocytes but consistent and robust increase in human astrocytes. Genes encoding additional MHC I-interacting antigen processing and presenting proteins, such as Tap1, Tap2, and ICAM1 showed similarly robust increases in human astrocytes but no change in mouse astrocytes treated with Poly I:C (Supplemental Table 7, 8).

To identify coregulated gene networks changing under Poly I:C or TNFα treatment, we performed WGCNA and identified a module (black) upregulated in human astrocytes, but not mouse astrocytes under both treatment conditions (Supplemental Table 6 and Supplemental Figure 4). This module is involved in inflammatory responses to double stranded RNAs. The network analyses corroborated the finding that, at the specific dosage of Poly I:C or TNFα we used, human astrocytes showed stronger inflammatory responses compared to mouse astrocytes.

Signaling pathways downstream of Poly I:C treatment have been well characterized in multiple cell types of the immune system. Poly I:C binds one of the pattern-recognition receptors that recognize “danger” signal, the Toll-like receptor 3 (Tlr3), located in endosomes. TLR3 signals through an adaptor protein, Myd88, which activates the NFκB signaling pathway (Rel, Relb, Nfkb1, Nfkb2, etc). NFκB activation and nuclear translocation in turn activates interferon signaling (interferon responsive genes include Irf1, Irf2, Ifr7, and Irf9). NFκB signaling pathway cross talks with Stat3, the phosphorylation of which is involved in astrocyte reactivity. We found that all of the above-mentioned molecules are strongly upregulated after Poly I:C treatment in human astrocytes but showed modest or no upregulation in mouse astrocytes (Figure 7F). We found a similar pattern of stronger activation of these genes in TNFα-treated human astrocytes compared to mouse astrocytes (Figure 7G).

### Poly I:C and TNF**α** induced common transcriptional responses distinct from hypoxia responses

Do different types of perturbations induce a shared core astrocyte reactivity program or distinct programs specific to each perturbation? We found that, in both species, very few genes are induced by all three stimuli: hypoxia, poly I:C, and TNFα (Supplemental Figure 7). Poly I:C and TNFα induced many common gene changes, but these genes are largely different from hypoxia induced genes. WGCNA results corroborated this finding (Supplemental Figure 4). Therefore, consistent with previous observations, our data supports the idea that astrocytes have a diverse toolbox to react to diseases and perturbations, and they selectively use a subset of the tools in a context-dependent manner (Khakh and Sofroniew, 2015; Sofroniew, 2005; Sofroniew and Vinters, 2010; Zamanian et al., 2012).

## Discussion

In this study, we identified the conservation and divergence of astrocytic responses to disease-relevant perturbations between human and mouse. Using methods established to isolate and culture resting/homeostatic astrocytes from developmentally matched human and mouse astrocytes in parallel, and from which stimulate these mouse and human astrocytes with equivalent, controlled experimental paradigms for direct comparison, we found important differences between both resting and reactive human and mouse astrocytes. We found that, at rest, mouse and human astrocytes have difference in their mitochondria respiration rates. We also found that in response to different stimuli, reactive astrocytes show species specific differences. Human astrocytes are more susceptible to oxidative stress than mouse astrocytes, potentially contributing to differences between mouse models and human patients with neurodegeneration. Furthermore, we found that hypoxia induces a pro-growth molecular program in mouse astrocytes but not human astrocytes, potentially underlying greater functional recovery in mouse models of ischemic stroke than human patients. In addition, we found that Poly I:C and TNFα induce antigen presenting genes in human but not mouse astrocytes.

### The conservation and divergence of human and mouse astrocytes and their implications for translational research

We found extensive conservation in gene expression levels between human and mouse astrocytes but divergent responses of human and mouse astrocytes to certain stimuli. The discovery of divergent responses across species will inform, rather than obviate the use of the mouse as a model system. For example, we found that human astrocytes express much lower levels of the detoxification enzyme Catalase and are more susceptible to oxidative stress than mouse astrocytes. Since oxidative stress is one of the pathological hallmarks of neurodegeneration, one could “humanize” mouse models by using mice with reduced level of Catalase (e.g. heterozygotes for the Catalase gene) in modeling neurodegeneration.

The conservation and divergence of hypoxia responses of human and mouse astrocytes can be informative for ischemic stroke research. We showed that hypoxia induced HIF1 pathway activation, increase in glycolysis, and stimulation of autophagy in both species. However, only in mouse astrocytes did hypoxia induce a pro-growth molecular program. Human astrocytes appear perfectly able to sense oxygen shortage and make adaptive changes but stopped short in activating the pro-growth program. Investigation of how signal transduction occurs in mouse astrocytes that link hypoxia to the neuronal growth genes may lead to a therapy that activates the pro-neuronal growth program in human astrocytes.

We observed increased defense response gene expression in human compared to mouse astrocytes that was not solely dependent on environmental factors as shown in our xenograft experiment. Consistent with species specific differences in immune response, we further identified an increased expression of MHC class I molecules in response to inflammatory stimuli in human astrocytes compared to mouse. The role of MHC class 1 molecules in the CNS has yet to be fully characterized but appears to participate in developmental synaptic pruning (Lee et al., 2014) as well as the adaptive immune response. Alterations in MHC expression are a consistent observation in neurodegenerative disorders including Alzheimer’s disease, where they may contribute to synaptic dysfunction and altered T cell recruitment (Gate et al., 2020). Our discovery of species differences in astrocyte immune signaling has important implications for mouse models of neurodegeneration and may guide opportunities to enhance mechanistic understanding where mouse models display limited toxicity to human disease mutations and pathologies.

### Species differences in energy metabolism in astrocytes and other cell types

A few genes have been found to be associated with the expansion of cerebral cortex in human evolution (Florio et al., 2015; Hansen et al., 2010; Kalebic et al., 2019; Long et al., 2018; Mekel-Bobrov et al., 2005; Wang et al., 2011), one of which encodes a protein targeted to mitochondria, implicating metabolic changes in human brain evolution (Namba et al., 2020). It is unclear whether the species differences in metabolism we found are specific to astrocytes. Transcriptome and developmental differences have been found between human and mouse brains (Hodge et al., 2019; Johnson et al., 2009; Kang et al., 2011; Miller et al., 2014). However, there are very few reported respiration rate comparisons of human and mouse cells. One study that compared the metabolism of human and mouse muscle cells reported mixed results (Jacobs et al., 2013). At the organism level, our results are consistent with the observation that smaller mammals typically have higher metabolic rates per unit body weight than larger mammals (Perlman, 2016).

Mitochondria and energy metabolism changes are important in the pathogenesis of many neurological disorders. For example, many genes associated with Parkinson’s disease risk are involved in mitochondria function (Billingsley et al., 2019). A large cohort of genes involved in metabolism are induced after traumatic brain injury (Arneson et al., 2018). Impairment of glycolysis-derived metabolites in astrocytes contributes to cognitive deficits in Alzheimer’s disease (Le Douce et al., 2020). Previously, mitochondria function and energy metabolism have not been compared directly between human and mouse for any cell type of the central nervous system, to the best of our knowledge. Our new discovery of mitochondria and energy metabolism differences between human and mouse cells should be taken into consideration in translational research.

### Potential limitations of the study

All *in vitro* experiments in this study were performed using developing human and mouse astrocytes. Therefore, we do not recommend extrapolation of our conclusions to adult and/or aging contexts without further investigation. Although it is important to directly compare adult human and mouse astrocytes, it is challenging to obtain large numbers of fresh healthy brain tissue donations from adults to carry out characterization of astrocytic responses to disease-relevant stimuli with sufficient statistical power. Nevertheless, we performed RNA-seq of astrocytes purified from healthy brain tissue donated from adults (Supplemental Table 5). Analysis of adult astrocyte RNA-seq data showed human-mouse divergent pathways consistent with our *in vitro* findings, suggesting potential species differences consistent between developing and adult stages (Figure 5A, B).

Astrocytes from different brain regions differ in their morphology, physiology, and molecular phenotypes (Chai et al., 2017b; Farmer et al., 2016; Glasgow et al., 2014; Hochstim et al., 2008; John Lin et al., 2017; Miller et al., 2019; Molofsky et al., 2014; Zhang and Barres, 2010). To make our cross-species comparison of astrocytes accurate, we used the same brain region, cerebral cortex, from humans and mice as much as we could. Within the same region, astrocytes may be also heterogenous (Bayraktar et al., 2020; Darmanis et al., 2015; John Lin et al., 2017). There are morphologically distinct subtypes of astrocytes in humans but not in mice (Oberheim et al., 2006, 2009). Future studies utilizing single cell-based approaches may be able to further dissect the subpopulation level conservation and divergence between human and mouse astrocytes.

## Acknowledgments

We thank Baljit Khakh, Mark Sharpley, Michael Sofroniew, Jill Haney, Ajit Divakaruni for advice. We thank the UCLA Mitochondria Core, the Center for AIDS Research Core, the Eli and Edythe Broad Center of Regenerative Medicine and Stem Cell Research, UCLA BioSequencing Core Facility for their services, Kory Hamane, Mahnaz Akhavan and Suhua Feng for their technical support. This work is supported by the Achievement Rewards for College Scientists foundation Los Angeles Founder Chapter to M. I. G. the Dr. Sheldon and Miriam G. Adelson Medical Research Foundation to S.A.G., H.I.K., and D.H.G., the NIH/NINDS R00NS089780, R01NS109025, National Center for Advancing Translational Science UCLA CTSI Grant UL1TR001881, UCLA Eli and Edythe Broad Center of Regenerative Medicine and Stem Cell Research Innovation Award, and the Friends of the Semel Institute for Neuroscience & Human Behavior Friends Scholar Award to Y. Z.

## Author Contributions

J. L. and Y. Z. conceived of the project and designed the experiments. J. L. performed all experiments except those noted below. L.P. performed xenograft experiments and RNA-seq of xenografted astrocytes. M.I.G. contributed to the generation of RNA-seq libraries. M.C.C., A.G.A., and M.H. optimized xenografting conditions and assisted the xenograft experiments under the supervision of H.I.K. W.G.P., J.E.R., and D.H.G. performed WGCNA and analysis of FC correlations across species. Y-W.C. and X.Y. performed mapping of xenografted RNA-seq reads to human and mouse genomes. L. S. performed Seahorse Respirometry and TMRE/MTG imaging experiments. A.Y.C. and I.B.W. procured tissue samples. S.A.G. developed the xenograft method and provided training for xenograft experiments. J.L. and Y.Z. analyzed the data and wrote the paper. All authors read the manuscript.

## STAR Methods

### LEAD CONTACT AND MATERIALS AVAILABILITY

Further information and requests for resources and reagents should be directed to and will be fulfilled by the Lead Contact, Ye Zhang. This study did not generate new unique reagents.

### EXPERIMENTAL MODEL AND SUBJECT DETAILS

#### Experimental animals

All animal experimental procedures were approved by the Chancellor’s Animal Research Committee at the University of California, Los Angeles and conducted in compliance with national and state laws and policies. We used C57BL6 mice group-housed in standard cages (2-3 per cage). Rooms were maintained on a 12-hour light/dark cycle. Euthanasia and preparation of primary cultures of astrocytes were performed during the light cycle. 8-10 mixed sex pups at P1-3 from one to two litters were combined to make each batch of astrocyte culture.

#### Human tissue samples

Fetal human brain tissue without identifiable personal information was obtained with informed consent following elective pregnancy termination with exemption determination from the UCLA Office of the Human Research Protection Program. Gestational week 17-20 brain tissue was immersed in 4°C Dulbecco’s Phosphate Buffered Saline (DPBS, Gibco, 14040182) and transferred to the lab for tissue dissociation. In cases with largely intact brain tissue, we used whole cerebral cortex for astrocyte purification. In cases with fragmented tissue, we used fragments most likely to be cerebral cortex (typically large thin sheets). We included both female and male brain tissue. Samples sizes were noted in figure legends for each experiment.

#### Primary cell culture

Primary astrocyte cultures from humans and mice were generated by immunopanning and maintained in humidified 37°C incubator with 10% CO_2_ (see Method Details below). Cells from both female and male were used.

### METHOD DETAILS

#### Immunopanning purification of astrocytes

To examine responses of human and mouse astrocytes to stressors, we purified human and mouse astrocytes from developmentally equivalent stages, to the best of our knowledge. In humans, astrocytogenesis starts during the second trimester and continues through the third trimester (Choi and Lapham, 1978; Elder and Major, 1988; Roessmann and Gambetti, 1986). Human astrocytes reach maturity roughly around one year of age as determined by gene expression (Johnson et al., 2009; Kang et al., 2011; Pletikos et al., 2014; Zhang et al., 2016), although their physiological and functional maturation timeline is unclear. In mouse, astrocytogenesis starts at the perinatal period (embryonic day 17.5 [E17.5] of a 19-day gestation) and peaks between postnatal day 0 and 14 (P0-P14) (Molofsky and Deneen, 2015). Mouse astrocytes reach maturity roughly around one month of age as determined by morphology and gene expression (Bushong et al., 2004; Zhang et al., 2014, 2016). A single cell RNA-seq study of human and mouse brains found that molecular features of gestational week 16-20 human brains are similar to postnatal day 0-5 (P0-P5) mouse brains (Zhong et al., 2020). Therefore, we purified astrocytes from gestational week 17-20 human brains and P1-3 mouse brains. Within the age range used in our study (gestational week 17-20 for human and P1-3 for mouse), we did not observe age-dependent differences in any of the assays we tested.

We started astrocyte purification experiments using human and mouse brain tissue within similar postmortem intervals. For human samples, we received tissue within 30 minutes to 1 hour postmortem. We then performed very simple dissection that took less than 3 minutes. For mouse samples, we needed to combine one to two litters of 8-10 mice to get enough cells for each experiment. Cerebral cortex dissection from all the mice typically took ∼45 minutes before we started astrocyte purification experiments. We purified human and mouse astrocytes according to a previously published immunopanning protocol (Li et al., 2019; Zhang et al., 2016). Briefly, we coated three 150 mm-diameter petri dishes first with species-specific secondary antibodies (see Key Resources Table) and then with an antibody against CD45 (BD550539, both human and mouse), a hybridoma supernatant against the O4 antigen (mouse) or an antibody against CD90 (BD550402, human), and an antibody against HepaCAM (R&D Systems, MAB4108), respectively. We dissected cerebral cortices from human and mouse in PBS and removed meninges. We then dissociated the tissue with 6 unit/ml papain at 34.5°C for 45 minutes. We then mechanically triturated the tissue with 5ml serological pipets in the presence of a trypsin inhibitor solution. We then depleted microglia/macrophages, oligodendrocyte precursor cells, and neurons from the single cell suspension by incubating the suspension sequentially on the CD45, O4 (for mouse), or CD90 (for human) antibody-coated petri dishes. We then incubated the single cell suspension on the HepaCAM antibody-coated petri dish. After washing away nonadherent cells with PBS, we lifted astrocytes bound to the HepaCAM antibody-coated petri dish using trypsin and plated them on poly-D-lysine coated plastic coverslips in a serum-free medium containing Dulbecco’s modified Eagle’s medium (DMEM) (LifeTechnologies 11960069), Neurobasal (LifeTechnologies 21103049), sodium pyruvate (LifeTechnologies 11360070), Sato (Foo et al., 2011), glutamine (LifeTechnologies 25030081), N-acetyl cysteine (Sigma A8199) and HBEGF (Sigma E4643). For most of H_2_O_2_, TNFα, hypoxia, and poly I:C treatment experiments with exceptions detailed below, astrocytes were plated on 24-well culture plates at 75-100k per well. Human and mouse astrocyte cultures have similar densities for every type of experiment. For high density cultures for H_2_O_2_ treatment, 30k astrocytes were plated in a 50ul droplet in the middle of pre-dried poly-D-lysine-coated plastic coverslips on 24-well plates. After settling down for 20 minutes at 37°C, additional media were added. For Seahorse Respiration Assays, astrocytes were plated at 100-250k/well density in Agilent Seahorse 96 well cell culture microplates (cat#101085-004). For TMRE/MTG imaging, astrocytes were plated at 25-50k/well density in dark-walled flat bottom 96 well assay plate (CORNING, cat#3603). For poly I:C treatment, astrocytes were plated directly on poly-D-lysine-coated 24-well culture plates (Fisher, cat#08-772-1) without coverslips because poly I:C addition often induced cells floating away from coverslips. To purify xenografted human astrocytes and host mouse astrocytes from adult host mouse brains, we dissociated whole brains using 20u/ml papain, depleted microglia/macrophages, oligodendrocytes, and oligodendrocyte precursor cells with anti-CD45 antibody, GalC hybridoma supernatant, and O4 hybridoma supernatant coated plates, respectively. Three consecutive plates with the same antibody were used for depletion of each cell type. We then collected astrocytes with anti-HepaCAM antibody coated plates.

#### Serum-selection purification of astrocytes

Human brain tissue was dissociated into single cell suspensions as described above and plated on poly-D-lysine coated 25 cm^2^ culture flasks (VWR, cat#10861-672) in DMEM (Gibco, cat#11960044) with 10% fetal bovine serum (Gibco, cat#16140071) and 2mM glutamine. After 4-6 days, we vigorously shake off cells in the top layer (neurons and other glia) and left astrocytes on the bottom layer. We then harvested astrocytes for RNA-seq.

#### RNA-seq

We purified total RNA using the miRNeasy Mini kit (Qiagen cat# 217004) and analyzed RNA concentration and integrity with TapeStation (Agilent) and Qubit. All samples have RNA integrity numbers higher than 8.4. We then generated cDNA using the Nugen Ovation V2 kit (Nugen), fragmented cDNAs using the Covaris sonicator, and generated sequencing libraries using the Next Ultra RNA Library Prep kit (New England Biolabs) with 9-10 cycles of PCR amplification. We sequenced the libraries with the Illumina HiSeq 4,000 sequencer and obtained standard 16.3□±□5.7 million (mean□±□standard deviation) 50□bp single end reads per sample.

#### RNA-seq data analysis

We mapped sequencing reads to human genome hg38 and mouse genome MM10 using the STAR package (Dobin et al., 2013) and HTSEQ (Anders et al., 2015) to obtain raw counts. We then used the EdgeR-Limma-Voom packages (Robinson et al., 2010) in R to obtain Reads per Kilobase per Million Mapped Reads (RPKM) values. We calculated differential gene expression with the DESeq2 package (Love et al., 2014). Statistical significance of the overlap between two groups of genes were determined using http://nemates.org/MA/progs/overlap_stats.cgi Significance of the difference between two correlation coefficients was calculated using http://vassarstats.net/rdiff.html?

#### Comparison of transcriptomes of acutely purified human and mouse astrocytes

We mapped RNA-seq data from our previously obtained acutely purified human and mouse astrocyte datasets (Zhang et al., 2016) as described above. The age of samples is described in Supplemental Table 5. We calculated percentile rankings of RPKMs of each gene in each human and mouse astrocyte sample. We excluded genes with maximal percentile ranking across all samples < 0.33 as these genes are not expressed or very lowly expressed in all samples. We then performed Welch’s T-test between human and mouse samples and post hoc multiple comparison adjustment using the FDR method (Benjamini and Hochberg, 1995). Genes with FDR<0.05 and human-mouse percentile ranking difference >0.4 were used for GO term and cellular component analyses using string-db.org (Szklarczyk et al., 2019). Test gene lists were compared to background gene lists including all genes expressed at RPKM>0.05 in astrocytes.

#### WGCNA

Expression values from human and mouse were merged into a single expression matrix using only one-to-one human-mouse orthologues. Genes were retained if they had >20% non-zero values and were subsequently log2 (+0.001) transformed. We removed expression variation unrelated to the effect of treatment using the linear regression model “expr ∼ (1 | replicate)”. This maintained differences within each replicate pair, capturing the effect of treatment, but regressed out differences between replicate pairs such as basal species differences, or technical differences such as sequencing batch. Network analysis was performed through WGCNA using biweight midcorrelation (bicor) to reduce sensitivity to outliers. A soft threshold power of 18 was chosen to achieve scale-free topology (r^2^>0.8). The topological overlap matrix was hierarchically clustered and modules defined using a minimum module size of 50 and deepSplit cut of 2. Module-trait correlations were used to assess whether a module was significantly associated with a particular treatment in a particular species.

#### H_2_O_2_ treatment

We treated human and mouse astrocytes cultured in 24-well plates with 100-500 μM H_2_O_2_ (Sigma cat#95321-100ML) and performed cell survival assay, Seahorse respiration assay, and mitochondria membrane potential assay described below.

#### Cell survival assay

We incubated human and mouse astrocytes with the live cell dye calcein-AM and the dead cell dye ethidium homodimer in LIVE/DEAD^TM^ Viability Kit (Invitrogen cat#L3224) for 10 minutes at room temperature protected from light and imaged the cells with an Evos FL Auto 2 inverted fluorescence microscope (Invitrogen) with a 10x lense.

#### Seahorse respiration assay

We cultured human and mouse astrocytes with media detailed above and changed to Seahorse assay medium with 10mM glucose, 2mM glutamine, 1mM pyruvate and 5mM HEPES on the day of Seahorse respiration assay. We used the Agilent Seahorse XFe96 Analyzer to measure oxygen concentration and extracellular pH changes. We first measured basal oxygen consumption rates at unperturbed conditions. We then added oligomycin to inhibit ATP-synthase (mitochondria Complex IV). The differences between basal and oligomycin conditions reflect the amount of oxygen consumption used for ATP production (Lardy et al., 1964). We next added carbonyl cyanide-4 (trifluoromethoxy) phenylhydrazone (FCCP), an uncoupling agent that collapses the proton gradient and disrupts the mitochondrial membrane potential (Park et al., 2002). As a result, electron flow through the electron transport chain is uninhibited, and oxygen consumption by complex IV reaches the maximum. Lastly, we added antimycin A to block Complex III and shut down mitochondria respiration (Ma et al., 2011). In the presence of antimycin A, measured respiration rate represents non-mitochondria respiration, with major contribution from the organelle, peroxisome (Reddy and Mannaerts, 1994). We made measurements every 5 minutes for 3-4 data points per condition. We sequentially added 2μM oligomycin, 0.5μM and 0.9μM FCCP, 2 μM antimycin A. After measurements, we stained cells with DAPI and counted the number of cells in each sample. Measurement results were then normalized by cell number.

#### Mitochondria membrane potential assay

We loaded cultured human and mouse astrocytes with 14nM TMRE and 200nM MTG and 1μg/ml Hoechst for 45 minutes, treated the cells with 100μM H_2_O_2_ and then measured fluorescence with imager at 1, 3, and 5 hours after H_2_O_2_ treatment. After staining, the cells were washed three times with culture medium containing 14nM TMRE to remove extra MTG and Hoechst dyes. TMRE and MTG fluorescence were imaged with an Operetta High-Content Imaging System (PerkinElmer). Fluorescence intensity after H_2_O_2_ treatment was normalized to untreated control.

#### Hypoxia treatment

We first cultured immunopanned human and mouse astrocytes under atmospheric oxygen concentrations for three days. We then cultured them under 1% oxygen for three days. Control cells were cultured under atmospheric oxygen concentrations for six days. We then harvested RNA for RNA-seq.

#### Poly I:C treatment

We first cultured immunopanned human and mouse astrocytes for three days. We then added 200ng/ml Poly I:C (Sigma, cat#P1530-25MG) to culture medium and cultured the cells for additional three days. We then harvested RNA for RNA-seq.

#### TNF**α** treatment

We treated human and mouse astrocytes 3 days in culture with 30ng/ml TNFα for 48 hours and harvested the cells for RNA-seq. We treated mouse astrocytes with TNFα from human (Cell Signaling Technology, 8902SF) and mouse sources (Cell Signaling Technology, 5178SF) and sequenced them in separate experiments. Similar number of genes were induced in mouse astrocytes by TNFα from human and mouse sources. We used cells treated with human TNFα for subsequent analyses.

#### Transplantation of human astrocytes into host mouse brains

We transplanted human astrocytes into host mouse brains according to published protocols (Benraiss et al., 2016; Han et al., 2013; Osipovitch et al., 2019; Wang et al., 2013; Windrem et al., 2008, 2014, 2017). Briefly, we purified human astrocytes as described above under the “Serum-selection purification of astrocytes” section. We then injected 100,000 cells per μl, 1 μl per injection, 4 injections per mouse at age P2-11. We used Rag2 immunodeficient mice to avoid graft rejection. The mice were maintained in autoclaved cages with autoclaved food and water in a specific pathogen-free facility.

#### Mapping xenograft reads to mouse to human genome using in silico combined human-mouse reference genome (ICRG)

RNA-seq data was assessed for quality parameters using FastQC (http://www.bioinformatics.babraham.ac.uk/projects/fastqc) and then trimmed with Trim_galore (https://www.bioinformatics.babraham.ac.uk/projects/trim_galore/). The RNA-seq reads were then mapped to ICRG using the method previously described (Callari et al., 2018). Briefly, reference genome and gene annotation files of human (hg38) and mouse (mm10) were downloaded from GENCODE (Frankish et al., 2018). Human chromosomes were tagged as “chr” whereas the mouse chromosomes were renamed as “m.chr”. The two fasta files for human and mouse were then concatenated and indexed using STAR aligner (Dobin et al., 2013), allowing only one top scored locus to be mapped if multiple mappings occur. Benchmarking results showed low false alignment rate in both pure human (0.74%) and pure mouse RNA-seq (2.74%). After alignment, bam files were separated based on the “chr” (human) and “m.chr” (mouse) labels, followed by read counting using Rsubread (Liao et al., 2019) to obtain the corresponding count matrix.

#### Non-supervised hierarchical clustering

We performed clustering in R using the hclust() function.

#### Immunohistochemistry

Mice were anesthetized with isoflurane and transcardially perfused with PBS followed by 4% PFA. Brains were removed and further fixed in 4% PFA at 4°C overnight. The brains were washed with PBS and cryoprotected in 30% sucrose at 4°C for two days before immersed in OCT (Fisher, cat#23-730-571) and stored at −80°C. Brains were sectioned on a cryostat (Leica) and 30μm floating sections were blocked and permeablized in 10% donkey serum with 0.2% Triton X-100 in PBS and then stained with primary antibodies against human nucleus protein (Chemicon, cat#MAB1281, Dilution 1:500) and human GFAP (Sternberger, cat#SMI21, Dilution 1:500) at 4°C overnight. Sections were washed three times with PBS and incubated for 2 hours at room temperature with secondary antibodies followed by three additional PBS washes. The sections were then mounted on Superfrost Plus micro slides (Fisher, cat#12-550-15) and covered with mounting medium (Fisher, cat#H1400NB) and glass coverslips.

### QUANTIFICATION AND STATISTICAL ANALYSIS

The number of patients, animals, or replicates is described in figures and figure legends. RNA-seq data were analyzed as described in detail in the RNA-seq section above. For all non-RNA-seq data and RNA-seq data comparison between species, analyses were conducted using R and the Prism 8 software (Graphpad). Normality of data was tested by the Shapiro-Wilke test. For data with normal distribution, Welch’s t test was used for two-group comparisons and one-way ANOVA was used for multi-group comparisons. For data that deviate from normal distribution, the Mann-Whitney test is used. Data from technical replicates from the same patient or the same litter of mice was averaged and used as a single biological replicate in statistical analyses. An estimate of variation in each group is indicated by the standard error of mean (S.E.M.). * p<0.05, ** p<0.01, *** p<0.001.

## Supplemental Information

### Supplemental Figures

**Supplemental Figure 1. Differences in percentile ranking of gene expression between serum-free cultured and acutely purified human astrocytes**

The numbers of genes that changed decile (>10% difference in percentile ranking) or stayed in the same decile (<10% difference) between the two conditions were shown. The majority (88.1%) of the genes are in the same decile in cultured and acutely purified astrocytes.

**Supplemental Figure 2. Correlation between gene expression of xenografted and acutely purified astrocytes.**

Spearman’s correlation coefficient ρ is shown in all panels.

(A) Acutely purified human astrocytes (all developing and adult ages) vs. transplanted human astrocytes. Only protein coding genes are included.

(B) Acutely purified mouse astrocytes (all developing and adult ages) vs. transplanted human astrocytes. Only protein coding genes are included.

(C) Acutely purified human fetal astrocytes vs. transplanted human astrocytes. Protein coding genes with average RPKM>0 are included.

(D) Acutely purified human children’s astrocytes vs. transplanted human astrocytes. Protein coding genes with average RPKM>0 are included.

(E) Acutely purified human adult astrocytes vs. transplanted human astrocytes. Protein coding genes with average RPKM>0 are included.

**Supplemental Figure 3. Survival of human and mouse astrocytes under oxidative stress.**

(A) Experimental design.

(B) Human and mouse astrocytes treated with H_2_O_2_ or medium control stained with the live cell dye calcein-AM (green) and the dead cell dye ethidium homodimer (red). Scar bar: 200μm.

(C) Survival rate.

(D) Survival rate of astrocytes treated with H_2_O_2_ normalized to the survival rate of medium control treated cells.

**Supplemental Figure 4. WGCNA identified gene coexpression modules**

(A) WGCNA dendrogram.

(B) Modules and their association with treatment conditions. The numbers on top in each rectangle represent correlation and the numbers on the bottom in parentheses are p-values.

**Supplemental Figure 5. Scatter plots and correlations of hypoxia, Poly I:C, and TNFα treatment-induced gene expression changes in human and mouse astrocytes**

Genes with average count <1 were filtered out.

**Supplemental Figure 6. Molecular responses of human and mouse astrocytes to Poly I:C or TNFα**

(A,B,D,E) Volcano plots of genes significantly different between Poly I:C or TNFα-treated and control conditions. Each red dot represents a significantly different gene.

(C, F) Comparison of fold change of Poly I:C or TNFα-induced genes in human and mouse astrocytes. Genes with FDR<0.05 in both species, FC>1.5, and average RPKM of control/treated >1 are included. Blue line: linear regression trendline. Black line: predicted trendline of y=x if there were no species differences.

(G) Antigen presentation-related GO terms enriched in TNFα-induced genes *only* in human astrocytes.

(H-M) Expression of top 3 highest expressing MHC Class I antigen presentation genes in TNFα-treated and control human and mouse astrocytes. n=6 litters of mice and 4 human patients.

**Supplemental Figure 7. Comparison of genes induced by hypoxia, Poly I:C, and TNFα treatment**

Protein coding genes with FDR<0.05, fold change >1.5 are included.

### Supplemental Tables

**Supplemental Table 1. Gene expression (RPKM) of acutely purified, serum-free culture, and serum-selected culture of human astrocytes.**

**Supplemental Table 2. GO terms associated with genes differentially expressed by serum-free cultured and acutely purified human astrocytes and hypoxia, Poly I:C, and TNF**α **treatment induced genes**

**Supplemental Table 3. GO terms associated with genes differentially expressed by human and mouse astrocytes.**

**Supplemental Table 4. Gene expression (RPKM) of grafted human astrocytes and host mouse astrocytes**

**Supplemental Table 5. Age of patients and mice used in RNA-seq comparison of astrocyte gene expression across species**

**Supplemental Table 6. WGCNA coexpression modules, member genes, and kMEs**

**Supplemental Table 7. Gene expression (RPKM) of hypoxia, Poly I:C, and TNF**α **treated astrocytes**

**Supplemental Table 8. Differentially expressed genes induced by hypoxia, Poly I:C, and TNF**α **treatment**

## References

Adams, K.L., and Gallo, V. (2018). The diversity and disparity of the glial scar. Nat. Neurosci. 21, 9–15.

Allen, N.J., Bennett, M.L., Foo, L.C., Wang, G.X., Chakraborty, C., Smith, S.J., and Barres, B.A. (2012). Astrocyte glypicans 4 and 6 promote formation of excitatory synapses via GluA1 AMPA receptors. Nature 486, 410–414.

Anders, S., Pyl, P.T., and Huber, W. (2015). HTSeq--a Python framework to work with high-throughput sequencing data. Bioinformatics 31, 166–169.

Anderson, M.A., Burda, J.E., Ren, Y., Ao, Y., O’Shea, T.M., Kawaguchi, R., Coppola, G., Khakh, B.S., Deming, T.J., and Sofroniew, M. V. (2016). Astrocyte scar formation aids central nervous system axon regeneration. Nature 532, 195–200.

Arneson, D., Zhang, G., Ying, Z., Zhuang, Y., Byun, H.R., Ahn, I.S., Gomez-Pinilla, F., and Yang, X. (2018). Single cell molecular alterations reveal target cells and pathways of concussive brain injury. Nat. Commun. 9, 3894.

Arnold, E.S., Ling, S.-C., Huelga, S.C., Lagier-Tourenne, C., Polymenidou, M., Ditsworth, D., Kordasiewicz, H.B., McAlonis-Downes, M., Platoshyn, O., Parone, P.A., et al. (2013). ALS-linked TDP-43 mutations produce aberrant RNA splicing and adult-onset motor neuron disease without aggregation or loss of nuclear TDP-43. Proc. Natl. Acad. Sci. U. S. A. 110, E736–45.

Bayraktar, O.A., Bartels, T., Holmqvist, S., Kleshchevnikov, V., Martirosyan, A., Polioudakis, D., Ben Haim, L., Young, A.M.H., Batiuk, M.Y., Prakash, K., et al. (2020). Astrocyte layers in the mammalian cerebral cortex revealed by a single-cell in situ transcriptomic map. Nat. Neurosci.

Benjamini, Y., and Hochberg, Y. (1995). Controlling the False Discovery Rate: A Practical and Powerful Approach to Multiple Testing. J. R. Stat. Soc. Ser. B 57, 289– 300.

Benraiss, A., Wang, S., Herrlinger, S., Li, X., Chandler-Militello, D., Mauceri, J., Burm, H.B., Toner, M., Osipovitch, M., Jim Xu, Q., et al. (2016). Human glia can both induce and rescue aspects of disease phenotype in Huntington disease. Nat. Commun. 7, 11758.

Bezard, E., Yue, Z., Kirik, D., and Spillantini, M.G. (2013). Animal models of Parkinson’s disease: Limits and relevance to neuroprotection studies. Mov. Disord. 28, 61–70.

Billingsley, K.J., Barbosa, I.A., Bandrés-Ciga, S., Quinn, J.P., Bubb, V.J., Deshpande, C., Botia, J.A., Reynolds, R.H., Zhang, D., Simpson, M.A., et al. (2019). Mitochondria function associated genes contribute to Parkinson’s Disease risk and later age at onset. Npj Park. Dis. 5, 8.

Blanco-Suarez, E., Liu, T.-F., Kopelevich, A., and Allen, N.J. (2018). Astrocyte-Secreted Chordin-like 1 Drives Synapse Maturation and Limits Plasticity by Increasing Synaptic GluA2 AMPA Receptors. Neuron 100, 1116–1132.e13.

Bordey, A., Lyons, S.A., Hablitz, J.J., and Sontheimer, H. (2001). Electrophysiological Characteristics of Reactive Astrocytes in Experimental Cortical Dysplasia. J. Neurophysiol. 85, 1719–1731.

Bozzo, F., Mirra, A., and Carrì, M.T. (2017). Oxidative stress and mitochondrial damage in the pathogenesis of ALS: New perspectives. Neurosci. Lett. 636, 3–8.

Bush, T.G., Puvanachandra, N., Horner, C.H., Polito, A., Ostenfeld, T., Svendsen, C.N., Mucke, L., Johnson, M.H., and Sofroniew, M. V (1999). Leukocyte infiltration, neuronal degeneration, and neurite outgrowth after ablation of scar-forming, reactive astrocytes in adult transgenic mice. Neuron 23, 297–308.

Bushong, E.A., Martone, M.E., and Ellisman, M.H. (2004). Maturation of astrocyte morphology and the establishment of astrocyte domains during postnatal hippocampal development. Int. J. Dev. Neurosci. 22, 73–86.

Callari, M., Batra, A.S., Batra, R.N., Sammut, S.-J., Greenwood, W., Clifford, H., Hercus, C., Chin, S.-F., Bruna, A., Rueda, O.M., et al. (2018). Computational approach to discriminate human and mouse sequences in patient-derived tumour xenografts. BMC Genomics 19, 19.

Chaboub, L.S., Manalo, J.M., Lee, H.K., Glasgow, S.M., Chen, F., Kawasaki, Y., Akiyama, T., Kuo, C.T., Creighton, C.J., Mohila, C.A., et al. (2016). Temporal Profiling of Astrocyte Precursors Reveals Parallel Roles for *Asef* during Development and after Injury. J. Neurosci. 36, 11904–11917.

Chai, H., Diaz-Castro, B., Shigetomi, E., Monte, E., Octeau, J.C., Yu, X., Cohn, W., Rajendran, P.S., Vondriska, T.M., Whitelegge, J.P., et al. (2017a). Neural Circuit-Specialized Astrocytes: Transcriptomic, Proteomic, Morphological, and Functional Evidence. Neuron 95, 531–549.e9.

Chai, H., Diaz-Castro, B., Shigetomi, E., Monte, E., Octeau, J.C., Yu, X., Cohn, W., Rajendran, P.S., Vondriska, T.M., Whitelegge, J.P., et al. (2017b). Neural Circuit-Specialized Astrocytes: Transcriptomic, Proteomic, Morphological, and Functional Evidence. Neuron 95, 531–549.e9.

Chen, Y., Miles, D.K., Hoang, T., Shi, J., Hurlock, E., Kernie, S.G., and Lu, Q.R. (2008). The basic helix-loop-helix transcription factor olig2 is critical for reactive astrocyte proliferation after cortical injury. J. Neurosci. 28, 10983–10989.

Choi, B.H., and Lapham, L.W. (1978). Radial glia in the human fetal cerebrum: A combined golgi, immunofluorescent and electron microscopic study. Brain Res. 148, 295–311.

Chung, W.-S., Clarke, L.E., Wang, G.X., Stafford, B.K., Sher, A., Chakraborty, C., Joung, J., Foo, L.C., Thompson, A., Chen, C., et al. (2013). Astrocytes mediate synapse elimination through MEGF10 and MERTK pathways. Nature 504, 394–400.

Chung, W.-S., Allen, N.J., and Eroglu, C. (2015). Astrocytes Control Synapse Formation, Function, and Elimination. Cold Spring Harb. Perspect. Biol. 7, a020370.

Crowley, L.C., Christensen, M.E., and Waterhouse, N.J. (2016). Measuring Mitochondrial Transmembrane Potential by TMRE Staining. Cold Spring Harb. Protoc. 2016, pdb.prot087361.

Darmanis, S., Sloan, S.A., Zhang, Y., Enge, M., Caneda, C., Shuer, L.M., Hayden Gephart, M.G., Barres, B.A., and Quake, S.R. (2015). A survey of human brain transcriptome diversity at the single cell level. Proc. Natl. Acad. Sci. 112, 7285–7290.

Dobin, A., Davis, C.A., Schlesinger, F., Drenkow, J., Zaleski, C., Jha, S., Batut, P., Chaisson, M., and Gingeras, T.R. (2013). STAR: ultrafast universal RNA-seq aligner. Bioinformatics 29, 15–21.

Le Douce, J., Maugard, M., Veran, J., Matos, M., Jégo, P., Vigneron, P.-A., Faivre, E., Toussay, X., Vandenberghe, M., Balbastre, Y., et al. (2020). Impairment of Glycolysis-Derived l-Serine Production in Astrocytes Contributes to Cognitive Deficits in Alzheimer’s Disease. Cell Metab. 31, 503–517.e8.

Dringen, R., Pawlowski, P.G., and Hirrlinger, J. (2005). Peroxide detoxification by brain cells. J. Neurosci. Res. 79, 157–165.

Elder, G.A., and Major, E.O. (1988). Early appearance of type II astrocytes in developing human fetal brain. Dev. Brain Res. 42, 146–150.

Eroglu, Ç., Allen, N.J., Susman, M.W., O’Rourke, N.A., Park, C.Y., Özkan, E., Chakraborty, C., Mulinyawe, S.B., Annis, D.S., Huberman, A.D., et al. (2009). Gabapentin Receptor α2δ-1 Is a Neuronal Thrombospondin Receptor Responsible for Excitatory CNS Synaptogenesis. Cell *139*, 380–392.

Farhy-Tselnicker, I., van Casteren, A.C.M., Lee, A., Chang, V.T., Aricescu, A.R., and Allen, N.J. (2017). Astrocyte-Secreted Glypican 4 Regulates Release of Neuronal Pentraxin 1 from Axons to Induce Functional Synapse Formation. Neuron 96, 428–445.e13.

Farmer, W.T., Abrahamsson, T., Chierzi, S., Lui, C., Zaelzer, C., Jones, E. V., Bally, B.P., Chen, G.G., Theroux, J.-F., Peng, J., et al. (2016). Neurons diversify astrocytes in the adult brain through sonic hedgehog signaling. Science (80-.). 351, 849–854.

Florio, M., Albert, M., Taverna, E., Namba, T., Brandl, H., Lewitus, E., Haffner, C., Sykes, A., Wong, F.K., Peters, J., et al. (2015). Human-specific gene ARHGAP11B promotes basal progenitor amplification and neocortex expansion. Science 347, 1465– 1470.

Foo, L.C., Allen, N.J., Bushong, E.A., Ventura, P.B., Chung, W.-S., Zhou, L., Cahoy, J.D., Daneman, R., Zong, H., Ellisman, M.H., et al. (2011). Development of a Method for the Purification and Culture of Rodent Astrocytes. Neuron 71, 799–811.

Frankish, A., Diekhans, M., Ferreira, A.-M., Johnson, R., Jungreis, I., Loveland, J., Mudge, J.M., Sisu, C., Wright, J., Armstrong, J., et al. (2018). GENCODE reference annotation for the human and mouse genomes. Nucleic Acids Res. 47, D766–D773.

Gandal, M.J., Haney, J.R., Parikshak, N.N., Leppa, V., Ramaswami, G., Hartl, C., Schork, A.J., Appadurai, V., Buil, A., Werge, T.M., et al. (2018). Shared molecular neuropathology across major psychiatric disorders parallels polygenic overlap. Science (80-.). 359, 693–697.

Gate, D., Saligrama, N., Leventhal, O., Yang, A.C., Unger, M.S., Middeldorp, J., Chen, K., Lehallier, B., Channappa, D., De Los Santos, M.B., et al. (2020). Clonally expanded CD8 T cells patrol the cerebrospinal fluid in Alzheimer’s disease. Nature 577, 399–404.

Glasgow, S.M., Zhu, W., Stolt, C.C., Huang, T.-W., Chen, F., LoTurco, J.J., Neul, J.L., Wegner, M., Mohila, C., and Deneen, B. (2014). Mutual antagonism between Sox10 and NFIA regulates diversification of glial lineages and glioma subtypes. Nat. Neurosci. 17, 1322–1329.

Han, X., Chen, M., Wang, F., Windrem, M., Wang, S., Shanz, S., Xu, Q., Oberheim, N.A., Bekar, L., Betstadt, S., et al. (2013). Forebrain engraftment by human glial progenitor cells enhances synaptic plasticity and learning in adult mice. Cell Stem Cell 12, 342–353.

Hansen, D. V., Lui, J.H., Parker, P.R.L., and Kriegstein, A.R. (2010). Neurogenic radial glia in the outer subventricular zone of human neocortex. Nature 464, 554–561.

Hay, M., Thomas, D.W., Craighead, J.L., Economides, C., and Rosenthal, J. (2014). Clinical development success rates for investigational drugs. Nat. Biotechnol. 32, 40–51.

Herrmann, J.E., Imura, T., Song, B., Qi, J., Ao, Y., Nguyen, T.K., Korsak, R.A., Takeda, K., Akira, S., and Sofroniew, M. V. (2008). STAT3 is a Critical Regulator of Astrogliosis and Scar Formation after Spinal Cord Injury. J. Neurosci. 28, 7231–7243.

Hochstim, C., Deneen, B., Lukaszewicz, A., Zhou, Q., and Anderson, D.J. (2008). Identification of Positionally Distinct Astrocyte Subtypes whose Identities Are Specified by a Homeodomain Code. Cell 133, 510–522.

Hodge, R.D., Bakken, T.E., Miller, J.A., Smith, K.A., Barkan, E.R., Graybuck, L.T., Close, J.L., Long, B., Johansen, N., Penn, O., et al. (2019). Conserved cell types with divergent features in human versus mouse cortex. Nature 573, 61–68.

Huang, Y.H., Sinha, S.R., Tanaka, K., Rothstein, J.D., and Bergles, D.E. (2004). Astrocyte glutamate transporters regulate metabotropic glutamate receptor-mediated excitation of hippocampal interneurons. J. Neurosci. 24, 4551–4559.

Ito, M., Aswendt, M., Lee, A.G., Ishizaka, S., Cao, Z., Wang, E.H., Levy, S.L., Smerin, D.L., McNab, J.A., Zeineh, M., et al. (2018). RNA-Sequencing Analysis Revealed a Distinct Motor Cortex Transcriptome in Spontaneously Recovered Mice After Stroke. Stroke 49, 2191–2199.

Jacobs, R.A., Díaz, V., Meinild, A., Gassmann, M., and Lundby, C. (2013). The C57Bl/6 mouse serves as a suitable model of human skeletal muscle mitochondrial function. Exp. Physiol. 98, 908–921.

John Lin, C.-C., Yu, K., Hatcher, A., Huang, T.-W., Lee, H.K., Carlson, J., Weston, M.C., Chen, F., Zhang, Y., Zhu, W., et al. (2017). Identification of diverse astrocyte populations and their malignant analogs. Nat. Neurosci. 20, 396–405.

Johnson, M.B., Kawasawa, Y.I., Mason, C.E., Krsnik, Ž., Coppola, G., Bogdanović, D., Geschwind, D.H., Mane, S.M., State, M.W., and Šestan, N. (2009). Functional and Evolutionary Insights into Human Brain Development through Global Transcriptome Analysis. Neuron 62, 494–509.

Kalebic, N., Gilardi, C., Stepien, B., Wilsch-Bräuninger, M., Long, K.R., Namba, T., Florio, M., Langen, B., Lombardot, B., Shevchenko, A., et al. (2019). Neocortical Expansion Due to Increased Proliferation of Basal Progenitors Is Linked to Changes in Their Morphology. Cell Stem Cell 24, 535–550.e9.

Kang, H.J., Kawasawa, Y.I., Cheng, F., Zhu, Y., Xu, X., Li, M., Sousa, A.M.M., Pletikos, M., Meyer, K.A., Sedmak, G., et al. (2011). Spatio-temporal transcriptome of the human brain. Nature 478, 483–489.

Kelley, K.W., Ben Haim, L., Schirmer, L., Tyzack, G.E., Tolman, M., Miller, J.G., Tsai, H.-H., Chang, S.M., Molofsky, A. V, Yang, Y., et al. (2018a). Kir4.1-Dependent Astrocyte-Fast Motor Neuron Interactions Are Required for Peak Strength. Neuron 98, 306–319.e7.

Kelley, K.W., Nakao-Inoue, H., Molofsky, A. V., and Oldham, M.C. (2018b). Variation among intact tissue samples reveals the core transcriptional features of human CNS cell classes. Nat. Neurosci. 21, 1171–1184.

Khakh, B.S., and Sofroniew, M. V (2015). Diversity of astrocyte functions and phenotypes in neural circuits. Nat. Neurosci. 18, 942–952.

Krencik, R., Weick, J.P., Liu, Y., Zhang, Z.-J., and Zhang, S.-C. (2011). Specification of transplantable astroglial subtypes from human pluripotent stem cells. Nat. Biotechnol. 29, 528–534.

Krencik, R., Hokanson, K.C., Narayan, A.R., Dvornik, J., Rooney, G.E., Rauen, K.A., Weiss, L.A., Rowitch, D.H., and Ullian, E.M. (2015). Dysregulation of astrocyte extracellular signaling in Costello syndrome. Sci. Transl. Med. 7, 286ra66.

Krencik, R., van Asperen, J. V., and Ullian, E.M. (2017a). Human astrocytes are distinct contributors to the complexity of synaptic function. Brain Res. Bull. 129, 66–73.

Krencik, R., Seo, K., van Asperen, J. V., Basu, N., Cvetkovic, C., Barlas, S., Chen, R., Ludwig, C., Wang, C., Ward, M.E., et al. (2017b). Systematic Three-Dimensional Coculture Rapidly Recapitulates Interactions between Human Neurons and Astrocytes. Stem Cell Reports 9, 1745–1753.

Lardy, H.A., Connelly, J.L., and Johnson, D. (1964). Antibiotics as Tools for Metabolic Studies. II. Inhibition of Phosphoryl Transfer in Mitochondria by Oligomycin and Aurovertin *. Biochemistry 3, 1961–1968.

Laug, D., Huang, T.-W., Huerta, N.A.B., Huang, A.Y.-S., Sardar, D., Ortiz-Guzman, J., Carlson, J.C., Arenkiel, B.R., Kuo, C.T., Mohila, C.A., et al. (2019). Nuclear factor I-A regulates diverse reactive astrocyte responses after CNS injury. J. Clin. Invest. 129, 4408–4418.

Lee, H., Brott, B.K., Kirkby, L.A., Adelson, J.D., Cheng, S., Feller, M.B., Datwani, A., and Shatz, C.J. (2014). Synapse elimination and learning rules co-regulated by MHC class i H2-Db. Nature.

Li, J., Khankan, R.R., Caneda, C., Godoy, M.I., Haney, M.S., Krawczyk, M.C., Bassik, M.C., Sloan, S.A., and Zhang, Y. (2019). Astrocyte-to-astrocyte contact and a positive feedback loop of growth factor signaling regulate astrocyte maturation. Glia.

Liao, Y., Smyth, G.K., and Shi, W. (2019). The R package Rsubread is easier, faster, cheaper and better for alignment and quantification of RNA sequencing reads. Nucleic Acids Res. 47, e47–e47.

Liddelow, S.A., and Barres, B.A. (2017). Reactive Astrocytes: Production, Function, and Therapeutic Potential. Immunity 46, 957–967.

Liddelow, S.A., Guttenplan, K.A., Clarke, L.E., Bennett, F.C., Bohlen, C.J., Schirmer, L., Bennett, M.L., Münch, A.E., Chung, W.-S., Peterson, T.C., et al. (2017). Neurotoxic reactive astrocytes are induced by activated microglia. Nature 541, 481–487.

Long, K.R., Newland, B., Florio, M., Kalebic, N., Langen, B., Kolterer, A., Wimberger, P., and Huttner, W.B. (2018). Extracellular Matrix Components HAPLN1, Lumican, and Collagen I Cause Hyaluronic Acid-Dependent Folding of the Developing Human Neocortex. Neuron 99, 702–719.e6.

Love, M.I., Huber, W., and Anders, S. (2014). Moderated estimation of fold change and dispersion for RNA-seq data with DESeq2. Genome Biol 15, 550.

Ma, X., Jin, M., Cai, Y., Xia, H., Long, K., Liu, J., Yu, Q., and Yuan, J. (2011). Mitochondrial electron transport chain complex III is required for antimycin A to inhibit autophagy. Chem. Biol. 18, 1474–1481.

Ma, Z., Stork, T., Bergles, D.E., and Freeman, M.R. (2016). Neuromodulators signal through astrocytes to alter neural circuit activity and behaviour. Nature 539, 428–432.

Majo, M., Koontz, M., Rowitch, D., and Ullian, E.M. (2020). An update on human astrocytes and their role in development and disease. Glia 68, 685–704.

Manwani, B., Liu, F., Xu, Y., Persky, R., Li, J., and McCullough, L.D. (2011). Functional recovery in aging mice after experimental stroke. Brain. Behav. Immun. 25, 1689–1700.

Masliah, E., Rockenstein, E., Veinbergs, I., Mallory, M., Hashimoto, M., Takeda, A., Sagara, Y., Sisk, A., and Mucke, L. (2000). Dopaminergic loss and inclusion body formation in alpha-synuclein mice: implications for neurodegenerative disorders. Science 287, 1265–1269.

McCarthy, K.D., and de Vellis, J. (1980). Preparation of separate astroglial and oligodendroglial cell cultures from rat cerebral tissue. J. Cell Biol. 85, 890–902.

Mekel-Bobrov, N., Gilbert, S.L., Evans, P.D., Vallender, E.J., Anderson, J.R., Hudson, R.R., Tishkoff, S.A., and Lahn, B.T. (2005). Ongoing adaptive evolution of ASPM, a brain size determinant in Homo sapiens. Science 309, 1720–1722.

Mergenthaler, P., Lindauer, U., Dienel, G.A., and Meisel, A. (2013). Sugar for the brain: the role of glucose in physiological and pathological brain function. Trends Neurosci. 36, 587–597.

Michalicová, A., Bhide, K., Bhide, M., and Kováč, A. (2017). How viruses infiltrate the central nervous system. Acta Virol. 61, 393–400.

Miller, J.A., Ding, S.-L., Sunkin, S.M., Smith, K.A., Ng, L., Szafer, A., Ebbert, A., Riley, Z.L., Royall, J.J., Aiona, K., et al. (2014). Transcriptional landscape of the prenatal human brain. Nature 508, 199–206.

Miller, S.J., Philips, T., Kim, N., Dastgheyb, R., Chen, Z., Hsieh, Y.-C., Daigle, J.G., Datta, M., Chew, J., Vidensky, S., et al. (2019). Molecularly defined cortical astroglia subpopulation modulates neurons via secretion of Norrin. Nat. Neurosci. 22, 741–752.

Minnerup, J., Sutherland, B.A., Buchan, A.M., and Kleinschnitz, C. (2012). Neuroprotection for stroke: current status and future perspectives. Int. J. Mol. Sci. 13, 11753–11772.

Molofsky, A.V., and Deneen, B. (2015). Astrocyte development: A Guide for the Perplexed. Glia 63, 1320–1329.

Molofsky, A. V., Kelley, K.W., Tsai, H.-H., Redmond, S.A., Chang, S.M., Madireddy, L., Chan, J.R., Baranzini, S.E., Ullian, E.M., and Rowitch, D.H. (2014). Astrocyte-encoded positional cues maintain sensorimotor circuit integrity. Nature 509, 189–194.

Molofsky, A. V, Krencik, R., Krenick, R., Ullian, E.M., Ullian, E., Tsai, H., Deneen, B., Richardson, W.D., Barres, B.A., and Rowitch, D.H. (2012). Astrocytes and disease: a neurodevelopmental perspective. Genes Dev. 26, 891–907.

Namba, T., Dóczi, J., Pinson, A., Xing, L., Kalebic, N., Wilsch-Bräuninger, M., Long, K.R., Vaid, S., Lauer, J., Bogdanova, A., et al. (2020). Human-Specific ARHGAP11B Acts in Mitochondria to Expand Neocortical Progenitors by Glutaminolysis. Neuron 105, 867–881.e9.

Nedergaard, M. (1994). Direct signaling from astrocytes to neurons in cultures of mammalian brain cells. Science 263, 1768–1771.

Nei, M., Xu, P., and Glazko, G. (2001). Estimation of divergence times from multiprotein sequences for a few mammalian species and several distantly related organisms. Proc. Natl. Acad. Sci. U. S. A. 98, 2497–2502.

Oberheim, N.A., Wang, X., Goldman, S., and Nedergaard, M. (2006). Astrocytic complexity distinguishes the human brain. Trends Neurosci. 29, 547–553.

Oberheim, N.A., Takano, T., Han, X., He, W., Lin, J.H.C., Wang, F., Xu, Q., Wyatt, J.D., Pilcher, W., Ojemann, J.G., et al. (2009). Uniquely hominid features of adult human astrocytes. J. Neurosci. 29, 3276–3287.

Oberheim Bush, N.A., and Nedergaard, M. (2017). Do Evolutionary Changes in Astrocytes Contribute to the Computational Power of the Hominid Brain? Neurochem. Res. 42, 2577–2587.

Osipovitch, M., Asenjo Martinez, A., Mariani, J.N., Cornwell, A., Dhaliwal, S., Zou, L., Chandler-Militello, D., Wang, S., Li, X., Benraiss, S.J., et al. (2019). Human ESC-Derived Chimeric Mouse Models of Huntington’s Disease Reveal Cell-Intrinsic Defects in Glial Progenitor Cell Differentiation. Cell Stem Cell 24, 107–122 e7.

Park, K.-S., Jo, I., Pak, K., Bae, S.-W., Rhim, H., Suh, S.-H., Park, J., Zhu, H., So, I., and Kim, K.W. (2002). FCCP depolarizes plasma membrane potential by activating proton and Na+ currents in bovine aortic endothelial cells. Pflugers Arch. 443, 344–352.

Parpura, V., Basarsky, T.A., Liu, F., Jeftinija, K., Jeftinija, S., and Haydon, P.G. (1994). Glutamate-mediated astrocyte–neuron signalling. Nature 369, 744–747.

Pascual, O., Casper, K.B., Kubera, C., Zhang, J., Revilla-Sanchez, R., Sul, J.-Y., Takano, H., Moss, S.J., McCarthy, K., and Haydon, P.G. (2005). Astrocytic purinergic signaling coordinates synaptic networks. Science 310, 113–116.

Patra, K.C., and Hay, N. (2014). The pentose phosphate pathway and cancer. Trends Biochem. Sci. 39, 347–354.

Perlman, R.L. (2016). Mouse models of human disease: An evolutionary perspective. Evol. Med. Public Heal. 2016, 170–176.

Pfrieger, F.W., and Barres, B.A. (1997). Synaptic efficacy enhanced by glial cells in vitro. Science 277, 1684–1687.

Pletikos, M., Sousa, A.M.M., Sedmak, G., Meyer, K.A., Zhu, Y., Cheng, F., Li, M., Kawasawa, Y.I., and Šestan, N. (2014). Temporal Specification and Bilaterality of Human Neocortical Topographic Gene Expression. Neuron 81, 321–332.

Puspita, L., Chung, S.Y., and Shim, J. (2017). Oxidative stress and cellular pathologies in Parkinson’s disease. Mol. Brain 10, 53.

Reddy, J.K., and Mannaerts, G.P. (1994). Peroxisomal Lipid Metabolism. Annu. Rev. Nutr. 14, 343–370.

Risher, W.C., Kim, N., Koh, S., Choi, J.-E., Mitev, P., Spence, E.F., Pilaz, L.-J., Wang, D., Feng, G., Silver, D.L., et al. (2018). Thrombospondin receptor α2δ-1 promotes synaptogenesis and spinogenesis via postsynaptic Rac1. J. Cell Biol. 217, 3747–3765.

Robel, S., Buckingham, S.C., Boni, J.L., Campbell, S.L., Danbolt, N.C., Riedemann, T., Sutor, B., and Sontheimer, H. (2015). Reactive astrogliosis causes the development of spontaneous seizures. J. Neurosci. 35, 3330–3345.

Robinson, M.D., McCarthy, D.J., and Smyth, G.K. (2010). edgeR: a Bioconductor package for differential expression analysis of digital gene expression data. Bioinformatics 26, 139–140.

Rodrigo, R., Fernandez-Gajardo, R., Gutierrez, R., Matamala, J., Carrasco, R., Miranda-Merchak, A., and Feuerhake, W. (2013). Oxidative Stress and Pathophysiology of Ischemic Stroke: Novel Therapeutic Opportunities. CNS Neurol. Disord. - Drug Targets 12, 698–714.

Rodriguez-Rodriguez, A., Egea-Guerrero, J., Murillo-Cabezas, F., and Carrillo-Vico, A. (2014). Oxidative Stress in Traumatic Brain Injury. Curr. Med. Chem. 21, 1201–1211.

Roessmann, U., and Gambetti, P. (1986). Astrocytes in the developing human brain. Acta Neuropathol. 70, 308–313.

Sasaguri, H., Nilsson, P., Hashimoto, S., Nagata, K., Saito, T., De Strooper, B., Hardy, J., Vassar, R., Winblad, B., and Saido, T.C. (2017). APP mouse models for Alzheimer’s disease preclinical studies. EMBO J. 36, 2473–2487.

Singh, S.K., Stogsdill, J.A., Pulimood, N.S., Dingsdale, H., Kim, Y.H., Pilaz, L.-J., Kim, I.H., Manhaes, A.C., Rodrigues, W.S., Pamukcu, A., et al. (2016). Astrocytes Assemble Thalamocortical Synapses by Bridging NRX1α and NL1 via Hevin. Cell 164, 183–196.

Sloan, S.A., Darmanis, S., Huber, N., Khan, T.A., Birey, F., Caneda, C., Reimer, R., Quake, S.R., Barres, B.A., and Paşca, S.P. (2017). Human Astrocyte Maturation Captured in 3D Cerebral Cortical Spheroids Derived from Pluripotent Stem Cells. Neuron 95, 779–790.e6.

Sofroniew, M. V. (2005). Reactive Astrocytes in Neural Repair and Protection. Neurosci. 11, 400–407.

Sofroniew, M. V (2014). Astrogliosis. Cold Spring Harb. Perspect. Biol. 7, a020420.

Sofroniew, M. V., and Vinters, H. V. (2010). Astrocytes: biology and pathology. Acta Neuropathol. 119, 7–35.

Springer, M.S., and Murphy, W.J. (2007). Mammalian evolution and biomedicine: new views from phylogeny. Biol. Rev. 82, 375–392.

Stogsdill, J.A., Ramirez, J., Liu, D., Kim, Y.H., Baldwin, K.T., Enustun, E., Ejikeme, T., Ji, R.-R., and Eroglu, C. (2017). Astrocytic neuroligins control astrocyte morphogenesis and synaptogenesis. Nature 551, 192–197.

Strack, A., Duffy, C.F., Malvey, M., and Arriaga, E.A. (2001). Individual Mitochondrion Characterization: A Comparison of Classical Assays to Capillary Electrophoresis with Laser-Induced Fluorescence Detection. Anal. Biochem. 294, 141–147.

Szklarczyk, D., Gable, A.L., Lyon, D., Junge, A., Wyder, S., Huerta-Cepas, J., Simonovic, M., Doncheva, N.T., Morris, J.H., Bork, P., et al. (2019). STRING v11: protein–protein association networks with increased coverage, supporting functional discovery in genome-wide experimental datasets. Nucleic Acids Res. 47, D607–D613.

Tasdemir-Yilmaz, O.E., and Freeman, M.R. (2014). Astrocytes engage unique molecular programs to engulf pruned neuronal debris from distinct subsets of neurons. Genes Dev. 28, 20–33.

Tchieu, J., Calder, E.L., Guttikonda, S.R., Gutzwiller, E.M., Aromolaran, K.A., Steinbeck, J.A., Goldstein, P.A., and Studer, L. (2019). NFIA is a gliogenic switch enabling rapid derivation of functional human astrocytes from pluripotent stem cells. Nat. Biotechnol. 37, 267–275.

Tönnies, E., and Trushina, E. (2017). Oxidative Stress, Synaptic Dysfunction, and Alzheimer’s Disease. J. Alzheimer’s Dis. 57, 1105–1121.

Tsai, H.-H., Li, H., Fuentealba, L.C., Molofsky, A. V., Taveira-Marques, R., Zhuang, H., Tenney, A., Murnen, A.T., Fancy, S.P.J., Merkle, F., et al. (2012). Regional Astrocyte Allocation Regulates CNS Synaptogenesis and Repair. Science (80-.). 337, 358–362.

Turrens, J.F. (2003). Mitochondrial formation of reactive oxygen species. J. Physiol. 552, 335–344.

Ullian, E.M., Sapperstein, S.K., Christopherson, K.S., and Barres, B.A. (2001). Control of Synapse Number by Glia. Science (80-.). 291, 657–661.

Vainchtein, I.D., Chin, G., Cho, F.S., Kelley, K.W., Miller, J.G., Chien, E.C., Liddelow, S.A., Nguyen, P.T., Nakao-Inoue, H., Dorman, L.C., et al. (2018). Astrocyte-derived interleukin-33 promotes microglial synapse engulfment and neural circuit development. Science (80-.). 359, 1269–1273.

Vakifahmetoglu-Norberg, H., Ouchida, A.T., and Norberg, E. (2017). The role of mitochondria in metabolism and cell death. Biochem. Biophys. Res. Commun. 482, 426–431.

Wang, S., Bates, J., Li, X., Schanz, S., Chandler-Militello, D., Levine, C., Maherali, N., Studer, L., Hochedlinger, K., Windrem, M., et al. (2013). Human iPSC-derived oligodendrocyte progenitor cells can myelinate and rescue a mouse model of congenital hypomyelination. Cell Stem Cell 12, 252–264.

Wang, X., Tsai, J.-W., LaMonica, B., and Kriegstein, A.R. (2011). A new subtype of progenitor cell in the mouse embryonic neocortex. Nat. Neurosci. 14, 555–561.

Windrem, M.S., Schanz, S.J., Guo, M., Tian, G.F., Washco, V., Stanwood, N., Rasband, M., Roy, N.S., Nedergaard, M., Havton, L.A., et al. (2008). Neonatal chimerization with human glial progenitor cells can both remyelinate and rescue the otherwise lethally hypomyelinated shiverer mouse. Cell Stem Cell 2, 553–565.

Windrem, M.S., Schanz, S.J., Morrow, C., Munir, J., Chandler-Militello, D., Wang, S., and Goldman, S.A. (2014). A competitive advantage by neonatally engrafted human glial progenitors yields mice whose brains are chimeric for human glia. J Neurosci 34, 16153–16161.

Windrem, M.S., Osipovitch, M., Liu, Z., Bates, J., Chandler-Militello, D., Zou, L., Munir, J., Schanz, S., McCoy, K., Miller, R.H., et al. (2017). Human iPSC Glial Mouse Chimeras Reveal Glial Contributions to Schizophrenia. Cell Stem Cell 21, 195–208.e6.

Xie, L., Kang, H., Xu, Q., Chen, M.J., Liao, Y., Thiyagarajan, M., O’Donnell, J., Christensen, D.J., Nicholson, C., Iliff, J.J., et al. (2013). Sleep drives metabolite clearance from the adult brain. Science 342, 373–377.

Yang, Y., Bazhin, A. V., Werner, J., and Karakhanova, S. (2013). Reactive Oxygen Species in the Immune System. Int. Rev. Immunol. 32, 249–270.

Yellen, G. (2018). Fueling thought: Management of glycolysis and oxidative phosphorylation in neuronal metabolism. J. Cell Biol. 217, 2235–2246.

Yu, X., Taylor, A.M.W., Nagai, J., Golshani, P., Evans, C.J., Coppola, G., and Khakh, B.S. (2018). Reducing Astrocyte Calcium Signaling In Vivo Alters Striatal Microcircuits and Causes Repetitive Behavior. Neuron 99, 1170–1187.e9.

Zamanian, J.L., Xu, L., Foo, L.C., Nouri, N., Zhou, L., Giffard, R.G., and Barres, B.A. (2012). Genomic analysis of reactive astrogliosis. J. Neurosci. 32, 6391–6410.

Zhang, Y., and Barres, B.A. (2010). Astrocyte heterogeneity: an underappreciated topic in neurobiology. Curr. Opin. Neurobiol. 20, 588–594.

Zhang, Y., and Barres, B.A. (2013). A smarter mouse with human astrocytes. Bioessays 35, 876–880.

Zhang, Y., Chen, K., Sloan, S.A., Bennett, M.L., Scholze, A.R., O’Keeffe, S., Phatnani, H.P., Guarnieri, P., Caneda, C., Ruderisch, N., et al. (2014). An RNA-Sequencing Transcriptome and Splicing Database of Glia, Neurons, and Vascular Cells of the Cerebral Cortex. J. Neurosci. 34, 11929–11947.

Zhang, Y., Sloan, S.A., Clarke, L.E., Caneda, C., Plaza, C.A., Blumenthal, P.D., Vogel, H., Steinberg, G.K., Edwards, M.S.B., Li, G., et al. (2016). Purification and Characterization of Progenitor and Mature Human Astrocytes Reveals Transcriptional and Functional Differences with Mouse. Neuron 89, 37–53.

Zheng, J., Winderickx, J., Franssens, V., and Liu, B. (2018). A Mitochondria-Associated Oxidative Stress Perspective on Huntington’s Disease. Front. Mol. Neurosci. 11, 329.

Zhong, S., Ding, W., Sun, L., Lu, Y., Dong, H., Fan, X., Liu, Z., Chen, R., Zhang, S., Ma, Q., et al. (2020). Decoding the development of the human hippocampus. Nature 577, 531–536.

