## Supplementary figures and images for "Conservation and divergence of vulnerability and responses to stressors between human and mouse astrocytes"

### Supplemental Figure 1

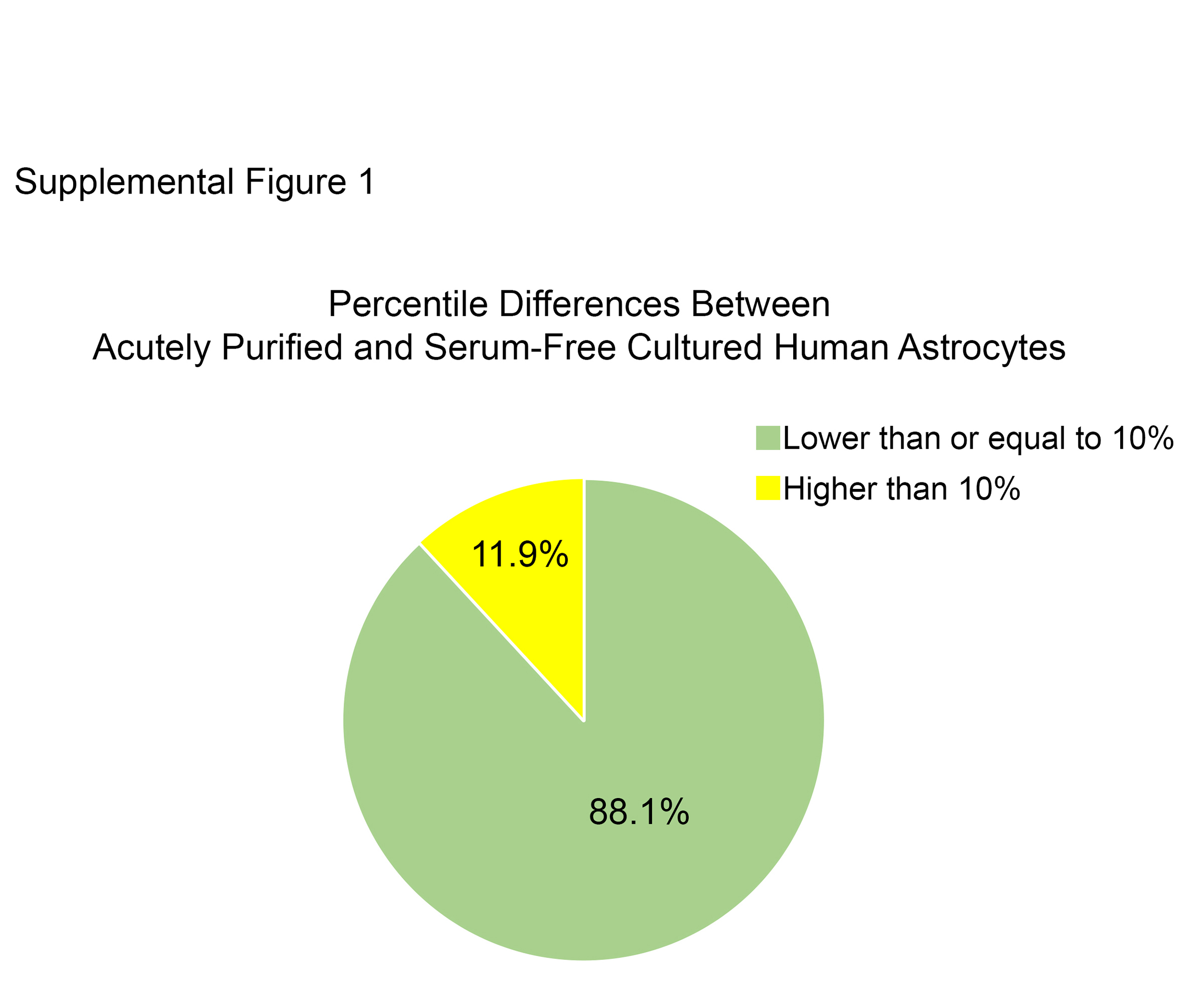

### Supplemental Figure 2

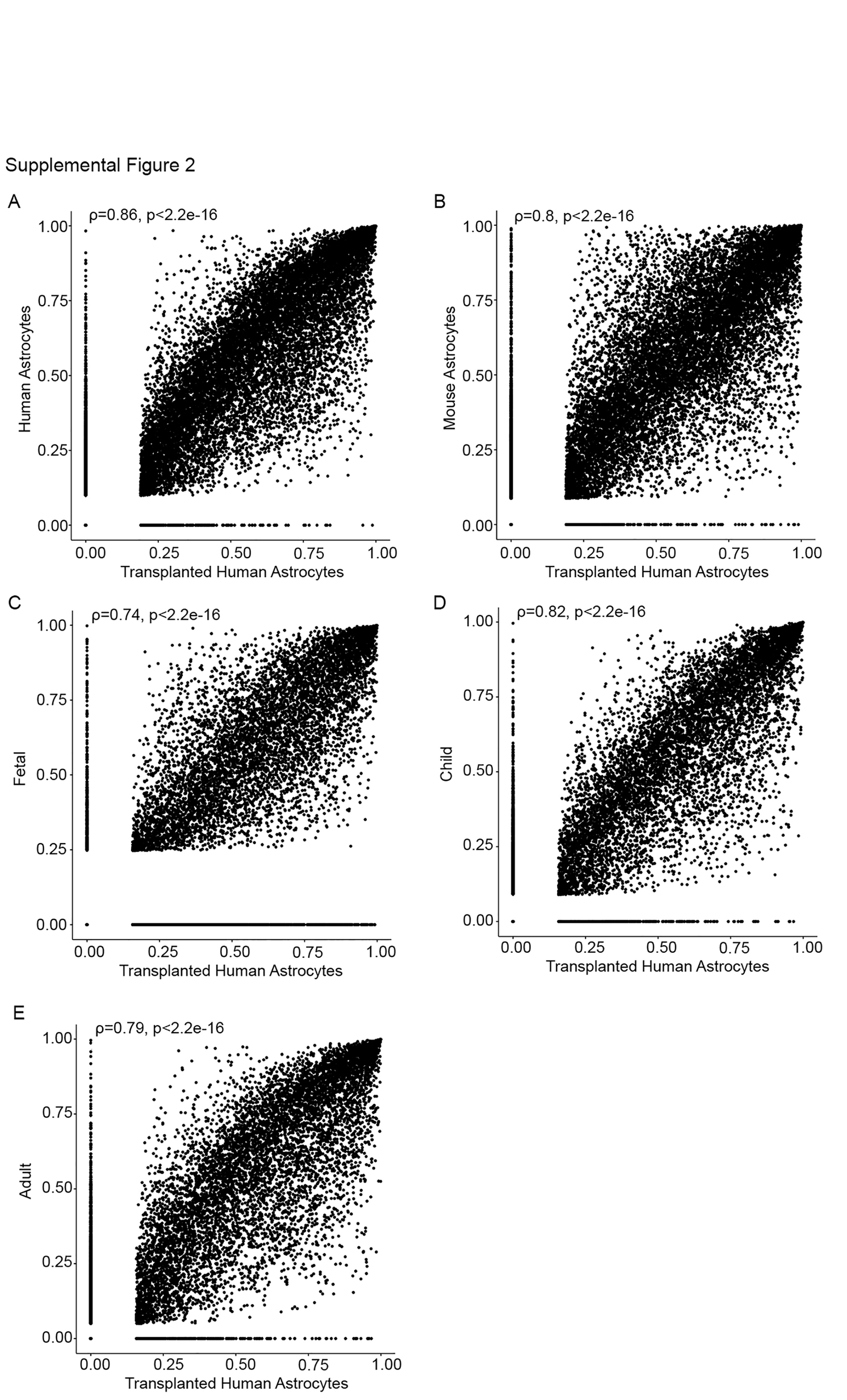

### Supplemental Figure 3

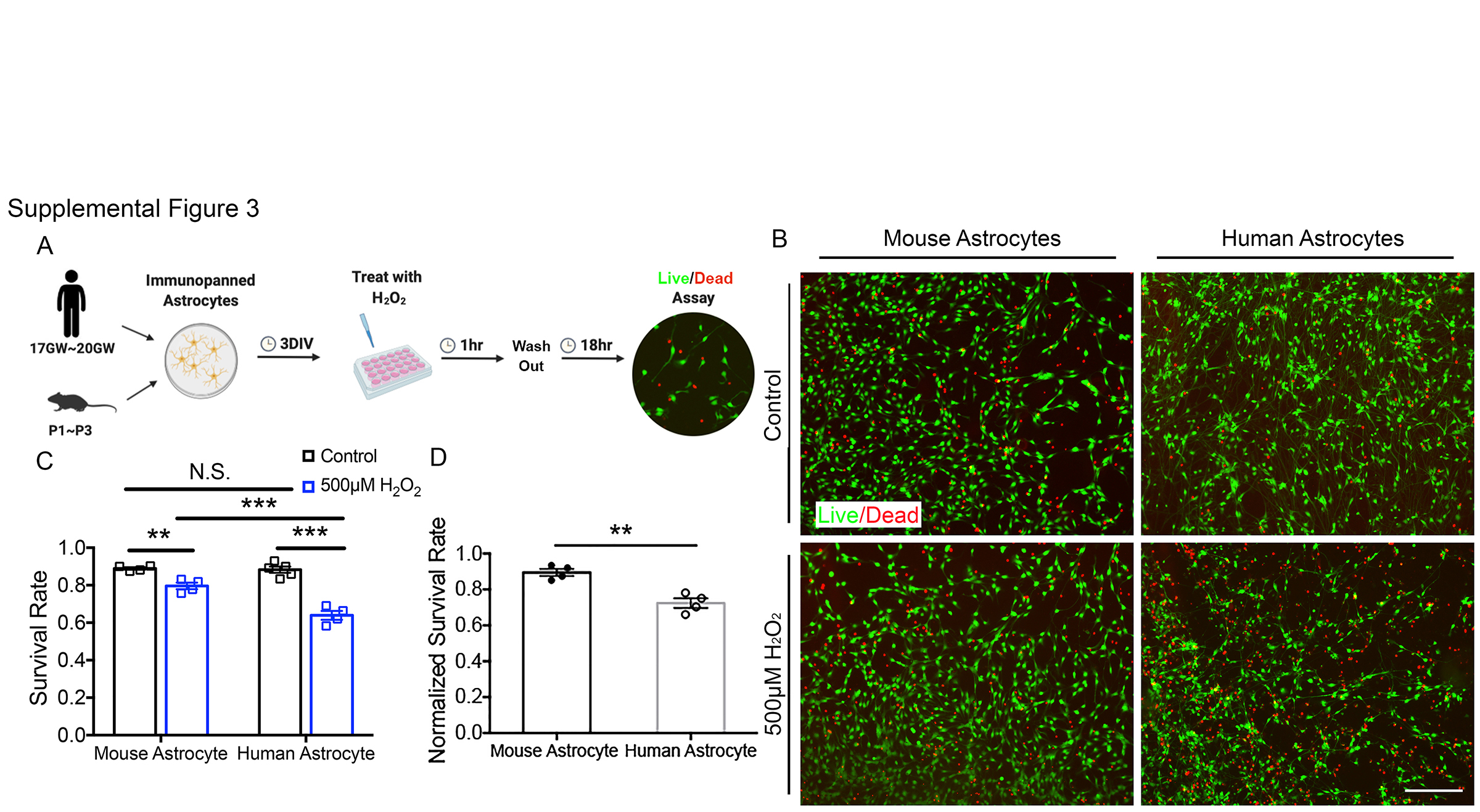

### Supplemental Figure 4

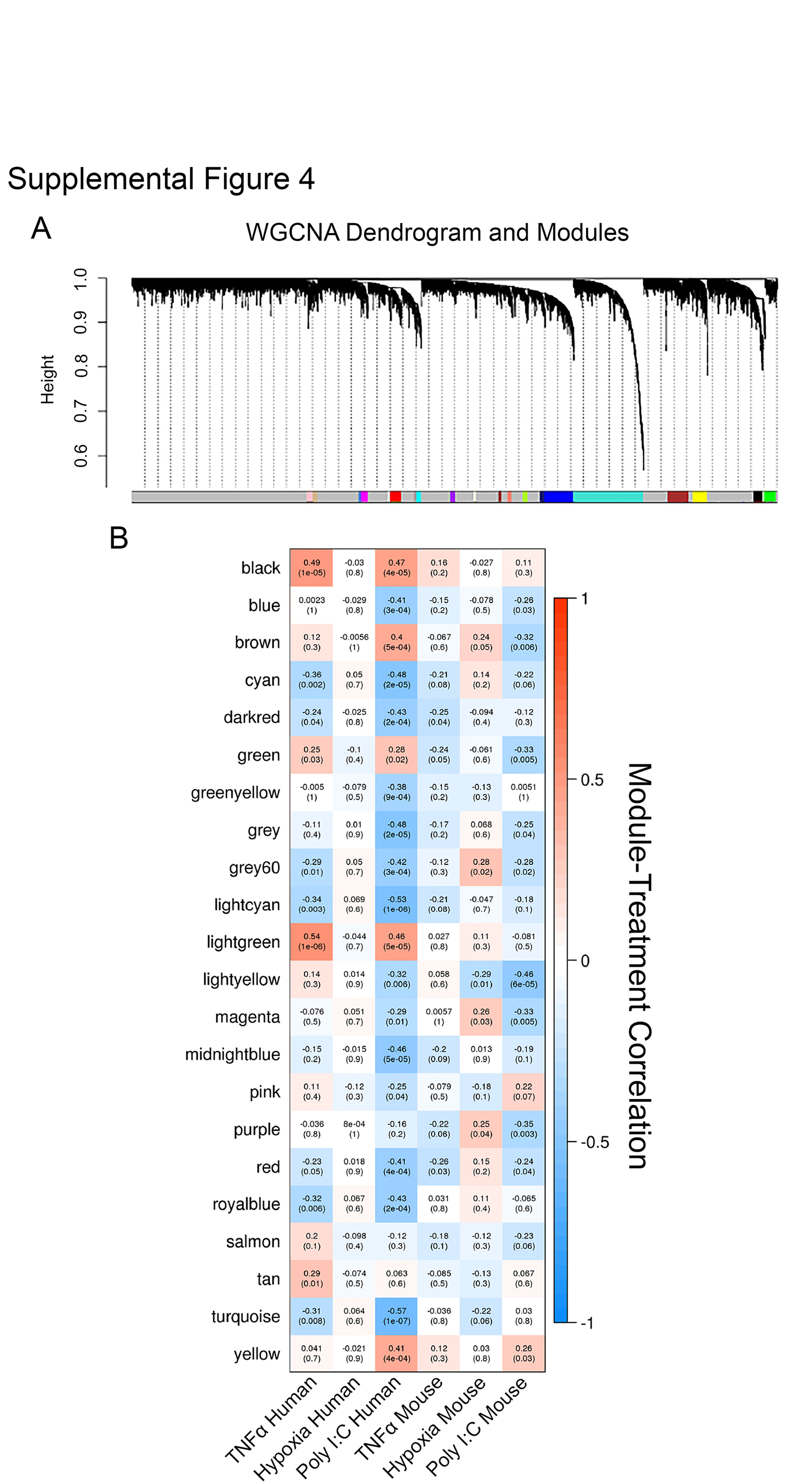

### Supplemental Figure 5

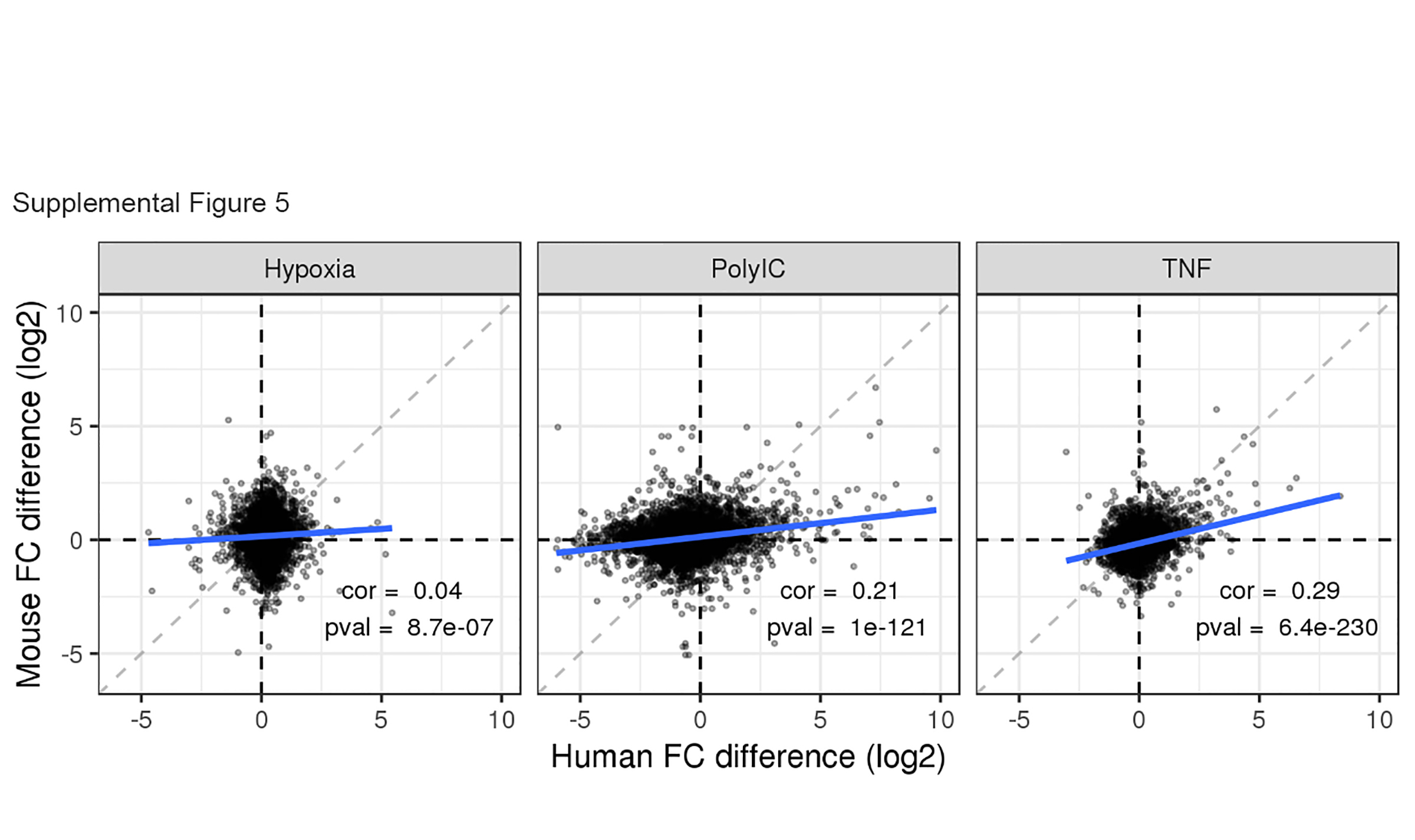

### Supplemental Figure 6

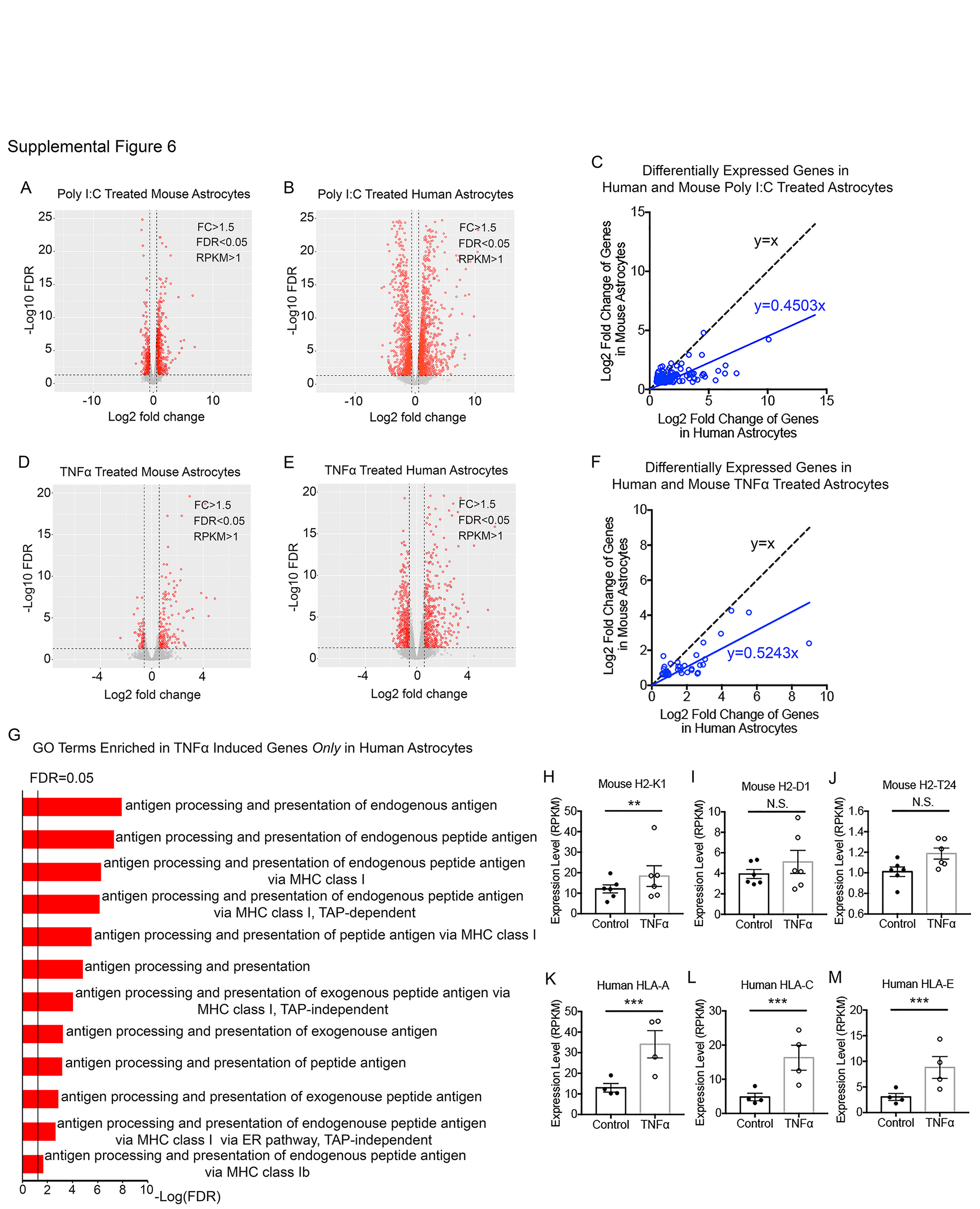

### Supplemental Figure 7

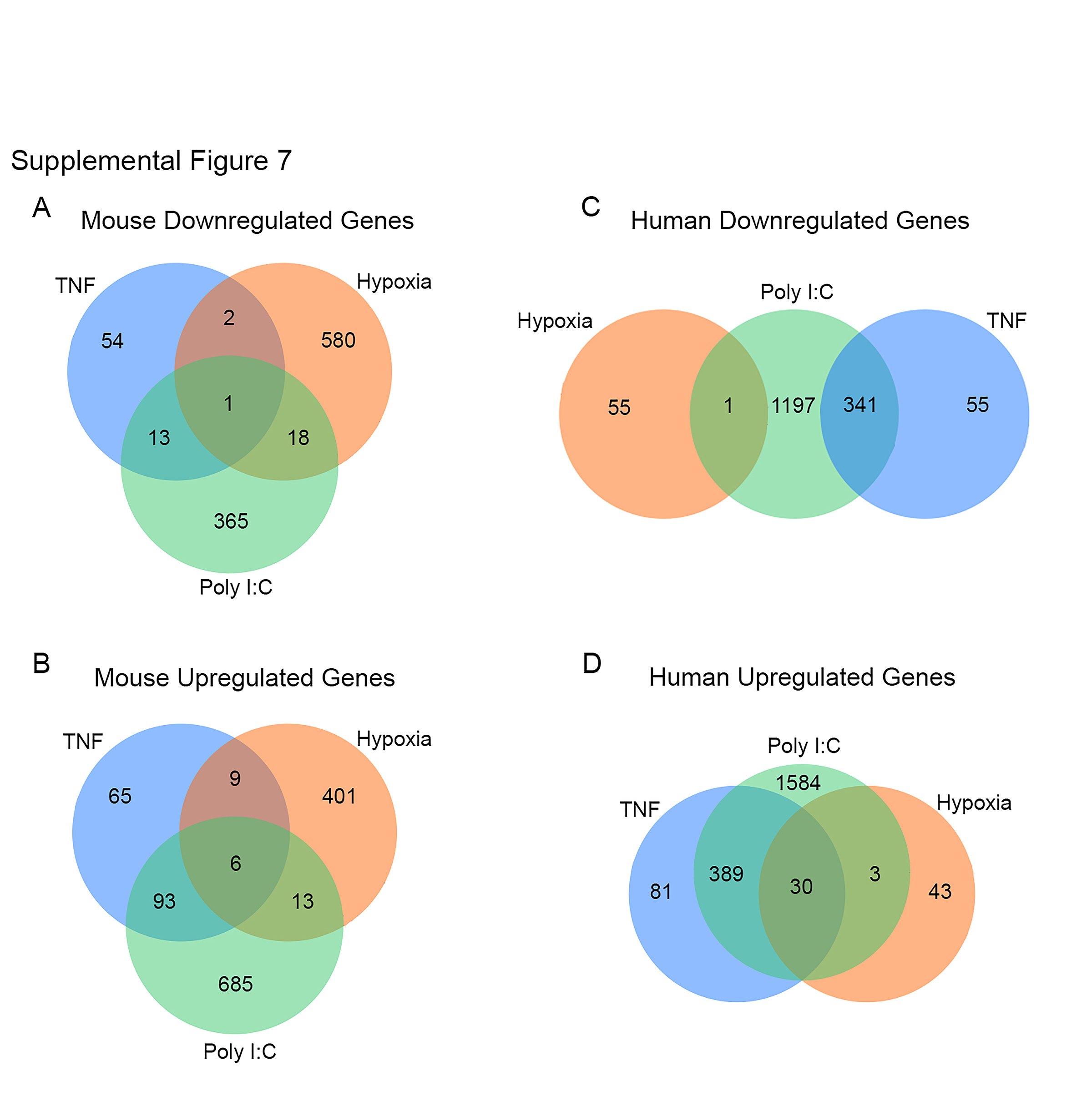
