## Supplemental Table 3 and 5 for "Conservation and divergence of vulnerability and responses to stressors between human and mouse astrocytes"

**Supplemental Table 3. Gene ontology (GO) terms enriched in genes differentially expressed by human and mouse astrocytes^a^**

| GO term | FDR |
| --- | --- |
| Higher in human |  |
| Defense response | 0.0042 |
| Higher in mouse |  |
| Cellular process | 0.0002 |
| Metabolic process  *Regulation of ph* | 0.0003 |
| Regulation of phosphorylation  R | 0.0023 |
| Regulation of phosphate metabolic process | 0.0035 |
| Regulation of catalytic activity | 0.0035 |
| Regulation of kinase activity | 0.0072 |
| Regulation of protein phosphorylation | 0.0088 |
| Negative regulation of cellular process | 0.0088 |
| Negative regulation of cellular metabolic process | 0.0094 |
| Regulation of protein modification process | 0.0094 |
| Cellular metabolic process | 0.0094 |
| Regulation of transferase activity | 0.0094 |
| Organic substance metabolic process | 0.0094 |
| Catabolic process | 0.0098 |
| Regulation of cellular protein metabolic process | 0,0110 |
| Nitrogen compound metabolic process | 0.0142 |
| Negative regulation of biological process | 0.0145 |
| Regulation of protein kinase activity | 0.0165 |
| Negative regulation of metabolic process | 0.0240 |
| Regulation of triglyceride biosynthetic process | 0.0240 |

^a^We used genes with FDR<0.05 and percentile ranking difference between species >0.4 for GO term analyses. Top 20 terms ranked by FDR (or all terms if there are fewer than 20 enriched terms) are shown.

**Supplemental Table 5. Age of patients and mice used in RNA-seq comparison of astrocyte gene expression across species**

| Patients/mice | Age |
| --- | --- |
| Human patients |  |
| 1 | 8 years |
| 2 | 13 years |
| 3 | 16 years |
| 4 | 21 years |
| 5 | 22 years |
| 6 | 35 years |
| 7 | 47 years |
| 8 | 51 years |
| 9 | 53 years |
| 10 | 60 years |
| 11 | 63 years |
| 12 | 63 years |
| Mice |  |
| Litter 1 | 7 days |
| Litter 2 | 7 days |
| Litter 3 | 1 month |
| Litter 4 | 4 months |
| Litter 5 | 7 months |
| Litter 6 | 9 months |
